# Harnessing Adaptive Chaos: Exploring the role of oxygen-ozone therapy in modulating complexity and repair in spine disorders via a bio-informatic model

**DOI:** 10.1101/2025.02.05.636618

**Authors:** Salvatore Chirumbolo, Marianno Franzini, Umberto Tirelli, Giovanni Ricevuti, Fortunato Loprete, Tommaso Richelmi, Francesco Vaiano, Antonio Carlo Galoforo, Marianna Chierchia, Luigi Valdenassi

## Abstract

Adaptive chaos represents a paradigm shift in understanding biological systems, where mild chaotic dynamics sustain homeostasis and resilience. This study delves into oxygen-ozone therapy’s ability to leverage adaptive chaos in treating spinal musculoskeletal disorders, particularly intervertebral disc degeneration (IVDD). Employing mathematical modelling and bioinformatic tools, the study evaluates chaos, complexity, and entropy across varying ozone doses to identify optimal therapeutic windows within a hormetic range. The results reveal that lower ozone doses (20–40 µg/ml O_3_) promote adaptive chaos, enhancing systemic complexity, reducing inflammatory cytokines, and fostering structural repair. Conversely, higher doses (60–80 µg/ml O_3_) induce excessive oxidative stress, exacerbating pathological chaos and impairing recovery. The integration of Shannon entropy, Lyapunov exponents, and fractal dimensions provides a novel framework for quantifying therapeutic outcomes. Findings highlight the role of controlled perturbations in stabilizing pathological systems, offering a dose-dependent roadmap for balancing flexibility and order. By correlating modelled insights with clinical evidence, the study underscores the significance of precision dosing, as the optimal range facilitates pain reduction and inflammatory control, correlating with improved complexity and reduced turbulence. This research pioneers the concept of adaptive chaos in medical interventions, emphasizing its translational potential for personalized and systemic medicine. Beyond IVDD, the findings establish a broader framework for adaptive chaos as a cornerstone of health restoration, advocating for therapies that optimize biological complexity. This work tries to bridge mathematical modelling and clinical evidence, offering a cutting-edge perspective on regenerative medicine and chaos-driven therapeutic strategies.

## Introduction

Adaptive chaos is an emerging paradigm that illuminates the intricate balance between stability and disorder in biological systems^1,2^. The concept of chaos in medicine is not particularly stressed, as the majority of biomedical phenomena and mechanisms are investigated by interpreting linear relationships between parts or at least networks integrating complex cross talking among different parts. Recent data outlines chaos as referring to the dynamic interplay where mild chaotic behaviour in one part of a system compensates for dysfunction in another, maintaining homeostasis and preventing catastrophic failure^2,3^. Contrary to conventional views that equate chaos with pathology, research reveals its crucial role in physiological flexibility and resilience. For instance, healthy cardiac rhythms exhibit chaotic properties^4,5^, enabling adaptation to stress, whereas excessively regular rhythms, such as those in heart arrhythmias, often indicate dysfunction^6,7^.

The principle of adaptive chaos provides a framework for understanding compensatory mechanisms in the body. As recently posited by Golbin and Umantsev^2^, mild disorders may stabilize broader systemic functions. For example, chaos in brain physiology reflects its dynamic adaptability, enabling rapid responses to stimuli. Neural chaos underlies complex functions like memory, perception, and coordination, with fractal patterns observed in EEGs^8^. While excessive chaos disrupts cognition, optimal levels balance order and flexibility, promoting resilience and efficient processing within neural networks^9^.

Beyond neurology, chaos plays a pivotal role also in immune responses.^10-12^

A controlled degree of inflammatory variability prevents extremes, either overreaction leading to autoimmunity or underreaction causing infection, thereby maintaining a responsive and balanced system, a model which originates from complex networking functions in immunity^13,14^.

In therapeutic contexts, adaptive chaos underscores the importance of controlled perturbations^2^. Oxygen-ozone therapy exemplifies this by leveraging oxidative stress in a hormetic manner^15-17^. Low doses of reactive oxygen species (ROS) stimulate repair and regulatory mechanisms, enhancing biological resilience and promoting healing^17,18^. This therapeutic strategy showcases how inducing controlled chaos can restore equilibrium in damaged tissues, offering a paradigm shift in medical treatment^16^. Traditional approaches that seek to suppress all chaotic symptoms may destabilize the system, eliminating compensatory mechanisms and exacerbating chronic conditions. Instead, therapeutic efforts should aim to optimize chaos within healthy bounds, fostering resilience and systemic stability^2,16^.

A sound example of ozone therapy is particularly promising in addressing symptomatic spine musculo-skeletal disorders, as it is able to face at the inflammatory concern related to this pathological disorder^19-23^.

Intervertebral disc degeneration (IVDD), a condition marked by chronic pain and reduced mobility, exemplifies the challenges posed by disrupted biological complexity^24-27^. Its pathophysiology involves a collapse of dynamic equilibrium, reflected in heightened inflammation, increased entropy, and impaired interactions between extracellular matrix (ECM) components and immune responses, including also a genetic landscape leading to IVDD^28^. Adaptive chaos offers a novel lens to understand and address these dynamics, particularly through oxygen-ozone therapy. This therapy modulates inflammation and immune responses via dose-dependent effects on cytokine profiles. For example, ozone stimulates anti-inflammatory cytokines like IL-10 while suppressing pro-inflammatory mediators such as TNF-α, fostering a transition from chaotic inflammation to structured healing^29,30^.

Despite its promise, the mechanisms by which ozone modulates chaos and complexity are underexplored. This study employs mathematical modelling to elucidate these effects, leveraging tools such as Ordinary Differential Equations (ODEs) and metrics like the Lyapunov exponent and Shannon entropy. Two scenarios, Case A (degeneration) and Case B (healing), simulate the biological transitions induced by ozone therapy. In Case A, high chaos reflects disordered inflammation, with elevated pro-inflammatory cytokines, immune cell activation, and ECM breakdown. Case B, by contrast, captures the resolution phase, characterized by reduced chaos, regulatory immunity, and enhanced repair.

Key variables in this model include cytokine levels, immune cell subsets, ECM degradation products, and bacterial contamination. Gaussian noise introduces biological variability, mirroring the chaotic instability of pathological states. By analyzing ozone’s impact on these variables, the study identifies optimal therapeutic windows within a hormetic range of concentrations (20–80 µg/ml). Lower doses, associated with reduced entropy and improved fractal dimensions, are hypothesized to enhance adaptive chaos and promote recovery. Higher doses, however, may exacerbate pathological chaos, highlighting the importance of precision in clinical protocols.

Metrics such as the Adaptive Chaos Metric (ACM) integrate entropy, complexity, and Lyapunov dynamics to quantify therapeutic outcomes. These modelled insights align with clinical data, correlating improvements in pain reduction and inflammatory markers with optimized ozone doses. This alignment bridges theoretical constructs with real-world outcomes, enhancing the translational relevance of adaptive chaos in therapeutic strategies.

The broader implications of adaptive chaos extend to personalized medicine, where treatments are tailored to preserve and enhance biological complexity. This perspective shifts the focus from suppressing symptoms to modulating systemic dynamics, fostering resilience and adaptability. For conditions like degenerative disc disease, this approach not only advances understanding of ozone’s mechanistic effects but also establishes a framework for evaluating adaptive responses in complex systems.

In conclusion, adaptive chaos is a cornerstone concept in modern medicine, offering insights into the reparative mechanisms of biological systems. By reframing mild chaotic phenomena as potentially beneficial, this paradigm enriches our understanding of health and disease. It advocates for therapies that enhance dynamic equilibrium, emphasizing the preservation of biological complexity as essential for effective medical care. The present study pioneers a multi-dimensional analysis of oxygen-ozone therapy’s impact on spinal disorders, bridging mathematical modelling with clinical evidence to optimize therapeutic strategies. This work not only deepens our grasp of ozone’s role in modulating chaos but also sets the stage for broader applications of adaptive chaos in personalized and systemic medicine.

## Materials and methods

### Modelling the system: simplified scenarios

Data modelling as a way to conceptualize and predict relationships in a system, even when data is incomplete or hypothetical was adopted. The resulting model should provide a framework to work with real data when they become available.

We considered two stages, in order to simplify our model.

Case A represents the intervertebral degenerated disc with herniation, before any successful therapeutical treatment involving oxygen-ozone, with the patient suffering from pain, disability and discomfort. This system is characterized by high inflammation, tissue breakdown, and immune activation. The dynamics include pro-inflammatory and anti-inflammatory cytokines, innate and adaptive immune cells, and ECM degradation products.

Key variables of Case A:

*I(t)* = pro-inflammatory cytokines (e.g., TNF-α, IL-1β, IL-6 and so forth), estimated as 100–500 pg/mL in local tissue or fluids, with TNFα around 50–200 pg/mL, central to pain and ECM breakdown, and IL-1β amounting to 10–50 pg/mL, driving matrix degradation.

*A(t)* = anti-inflammatory cytokines (e.g., IL-10, TGF-β), where the estimated concentration would be 10–50 pg/mL, notoriously less prominent than pro-inflammatory cytokines during active degeneration.

*M(t)* = innate immune cells (e.g., macrophages, neutrophils, dendritic cells, T and B cells, and so on), estimated as 10,000–100,000 cells/ml of disc tissue, represented by macrophages, the most abundant, predominantly M1 phenotype, by neutrophils, prominent in early stages but declining quickly, mast cells, present in smaller numbers (1–5% of total innate cells).

*T(t)* = T cells (e.g., CD4^+^Treg cells), intervertebral discs are typically sterile unless infected, estimating T cells (CD4^+^, CD8^+^, CD4^+^/CD25^+^/Foxp3^+^Tregs), around 1,000–10,000 cells/ml of tissue, where CD4^+^ T-cells being probably the dominant subset, amplifying inflammation, whereas CD25^+^Foxp3^+^Tregs less abundant (~5–10% of T-cell population). In this parameter, it has to be considered B cells, estimated as <1,000 cells/ml, typically low but present if there is an autoimmune or prolonged inflammatory response, and Natural Killer (NK) cells (500–2,000 cells/ml), present in small numbers, primarily targeting stressed or apoptotic disc cells.

*E(t)* = ECM degradation products (e.g., proteoglycans, collagen fragments, fibronectin fragments and aggrecan), estimated as 10–100 μg/mL, with aggrecan fragments around ~50–80 μg/mL in degenerated discs.

*B(t)* = bacteria concentration, estimated in ~10^3^–10^4^ colony-forming units (CFUs) in discs with infection (e.g., *Herbaspirillum, Ralstonia*, and *Burkolderia*).

Case A can be therefore modelled via the Ordered Differential Equations (ODE) operator as:

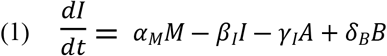

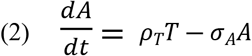

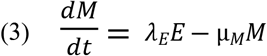

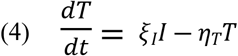

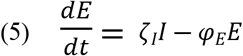

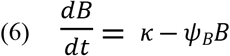

Where Greek letters are constant dependent from the different biological system evaluated.

Case B, reflects a resolved inflammation state with tissue repair and a shift toward regulatory immunity (healed patient, coming back to its dynamic equilibrium or health).

Key variables are *I(t), A(t), M(t), T(t)* and *E(t)*, as above but with significant differential estimations, *R(t)* is the repair signals (e.g., growth factors from M2 macrophages, promotion of collagen synthesis, and so on). Parameters estimated for case B are: a) innate immune cells (macrophages, mast cells, dendritic cells): 1,000–10,000 cells/ml, macrophages, as predominantly M2 phenotype (~80% of innate immune cells); mast cells with minimal activity (~1% of innate immune cells); b) inflammatory cytokines (TNF-α, IL-1β, IL-6): <10 pg/mL, barely detectable, reflecting resolved inflammation; c) anti-inflammatory cytokines (IL-10, TGF-β): 10–50 pg/mL, i.e., elevated compared to inflammatory cytokines, promoting tissue repair and fibrosis; d) molecules from damaged ECM (aggrecan, collagen fragments): <10 μg/mL, residual fragments are present but largely cleared during the resolution phase; e) bacteria: none detectable (sterile); f) T Cells (CD4^+^, CD8^+^, CD4^+^/CD25^+^Foxp3^+^Tregs): 500–1,000 cells/ml, where Tregs are in higher proportion (~20–30%) compared to degenerated discs, reflecting immune regulation; g) B Cells: <500 cells/ml, which means they are rarely present in a healed disc; h) Natural Killer (NK) Cells: 200–500 cells/ml, indicating minimal activity, reflecting the absence of apoptotic signals.

ODE modelling for case B:

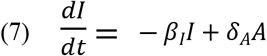

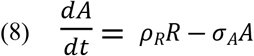

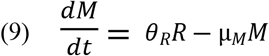

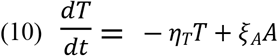

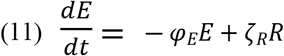

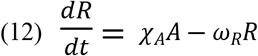

where Greek letters are constant dependent from the different biological system evaluated. Model simulation was performed taking into account the following issues.

Noise simulation was performed by adding Gaussian noise to each equation, where it can be supposed high noise in Case A (*σ*_*A*_ = 0.1) and lower noise in case B (*σ*_*B*_ = 0.01), then equations can be updated with noise:

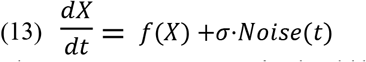

The Lyapunov exponent (λ) should be estimated as follows: a) positive λ = chaotic behaviour; b) zero λ or close to zero λ = low chaos, borderline chaos or periodic behaviour (adaptive chaos); c) negative λ close to zero = stable system; d) negative λ = de-complexed system.

### Modelling the system: chaos and complexity (part one)

We started with the concept that complexity is determined by the presence of numerous diverse components interconnected in functional and/or structural terms, for which scenario Shannon’s entropy, despite the large number of components, decreases, as this complexity is highly informed and informative. Conversely, the high number of free components, with the potential to establish relationships and thus generate information, represents a complexity with high informational (Shannon) entropy, which occurs when the system loses stability and enters a state of high chaos or turbulence. In such cases, the system may reduce the number of possible components to decrease chaotic instability, in order to rescue its dynamic equilibrium known as health.

Therefore, ordered differential equations were modelled according to the following pathways: a) high complexity arises from interconnected, functional, and structural relationships among components. This leads to low Shannon entropy because the system is well-informed and organized; b) low complexity occurs when components are free, unstructured, and unconnected, (for example molecules and organelles derived from tissue damages) leading to high Shannon entropy due to the system’s potential to generate relationships but failing to do so in a chaotic state; c) high chaos (instability) is reflected by a positive Lyapunov exponent. It corresponds to high Shannon entropy and low complexity due to disordered dynamics; d) stability (adaptive order) occurs with a very low positive or even negative Lyapunov exponent. It reduces Shannon entropy as relationships among components become more structured and informative.

For Case A, the simplified dynamic of pro-inflammatory cytokines is given by Eq. (1) where the production (+) was activated by innate immune cells (*M*) and bacteria (*B*), whereas reduction (−) occurs by neutralization via anti-inflammatory cytokines (A) and natural decay. The dynamic of anti-inflammatory cytokines is given by Eq. (2) (production stimulated by T cells (T) and decay occurring for natural degradation). Innate immune cells are given by ODE solutions of Eq. (3) (recruitment (+) as triggered by ECM degradation products (*E*), and decay (−), reflecting natural cell death). The dynamic of T cells is given by Eq. (4) (activation (+), is driven by pro-inflammatory cytokines (*I*), whereas decay (−), as the natural decline). The dynamic of ECM degradation products is given by Eq. (5) (generation (+) is stimulated by pro-inflammatory cytokines (*I*), whereas the clearance (−) reflects degradation product removal). Finally, bacteria dynamic is given by Eq. (6) (production (+), which describes external infection source, while clearance (−), represents immune response and treatment.

For Case B, the dynamic of pro-inflammatory cytokines is due to Eq. (7) (reduction (−), which reflects resolved inflammation caused by oxygen-ozone therapy and natural decay, and amplification (+), referring to the small contribution from anti-inflammatory cytokines (*A*)). The dynamic of anti-inflammatory cytokines is described by Eq. (8) (production (+), is driven by repair signals (*R*), whereas decay (−), via natural degradation), the one for innate immune cells from Eq. (9) (recruitment (+), triggered by repair signals (*R*), whereas decay (−), reflects natural decline, dynamic of T cells by Eq. (10) (reduction (−), reflects quiescence in the healed state and activation (+), driven by anti-inflammatory cytokines (*A*).

Finally, the dynamic of ECM degradation products is given by Eq. (11) (clearance (−), where ECM fragments are removed as healing progresses and regeneration (+), which is triggered by repair signals (*R*) and the dynamic of repair signals by Eq. (12) (production (+), stimulated by anti-inflammatory cytokines (*A*) and decay (−), representing natural dissipation.

The equations represent the interactions between inflammatory signals, immune cells, ECM components, and bacteria. They capture the dynamic balance of inflammatory and anti-inflammatory processes, ECM degradation, and repair. Case A includes mechanisms of inflammation, bacterial presence, and ECM degradation. Case B emphasizes repair and immune regulation, with lower inflammation and bacterial clearance. Furthermore, Gaussian noise was added to simulate physiological variability. High noise in Case A (before therapy) reflects chaotic, unregulated inflammation. Low noise in Case B (after therapy) reflects a more stable, adaptive system. The ODEs were solved using *scipy*.*integrate*.*solve_ivp* with the Runge-Kutta method. Gaussian noise (σ×randn()) was applied to simulate stochasticity. The Lyapunov exponent was calculated to quantify chaos by observing the divergence of nearby trajectories. Shannon entropy was used to measure system disorder, computed from the state probability distribution over time.

### Modelling the system: chaos and complexity (part two)

The systems for Case A (before therapy) and Case B (following therapy) can be also described using the following sets of coupled ordinary differential equations (ODEs). These equations capture the dynamics of immune cells, cytokines, and extracellular matrix (ECM) damage with the inclusion of noise to reflect biological variability.

In the Case A we could simplify the system as follows:

*I* = innate immune cells; *C*_*inflam*_ = pro-inflammatory cytokines; *C*_*anti*_ = anti-inflammatory cytokines; *E* = damaged ECM molecules; *η(t)* = noise term simulating biological variability. ODEs for Case A are given by:

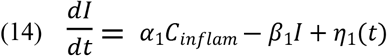

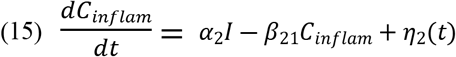

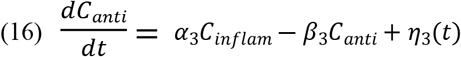

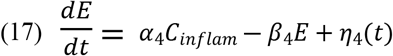

where, α_1_, α_2_, α_3_ and α_4_ > 0, indicating positive feedback promoting activation, the terms β_1_, β_2_, β_3_ and β_4_ > 0, indicating decay or regulatory terms reducing levels, and *η*_*i*_*(t) ~ N* (0, σ^2^), indicating Gaussian noise with high variance (σ^2^). This system exhibits chaotic dynamics due to the high variability (η_i_(t)) and strong positive feedback loops between inflammation and ECM damage.

For the healed disc (Case B), the system reflects reduced inflammation, low noise, and dominance of regulatory and reparative pathways. The variables remain the same, but the dynamics shift due to a predominance of anti-inflammatory cytokines and reparative processes.

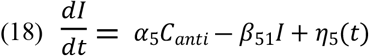

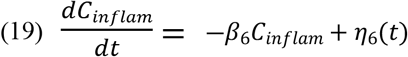

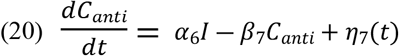

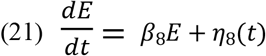

where α_5_ and α_6_ > 0, indicating repair-promoting factors, β_5_, β_6_, β_7_ and β_8_ > 0, representing strong decay terms reflecting stability, and *η_i_(t) ~ N* (0, σ^2^), indicating Gaussian noise with low variance (σ^2^).

This system is stable, with negative feedback loops leading to resolution of inflammation and ECM repair.

In order to evaluate the correct values of chaos dynamics and complexity, considering the system in its totality, the Lyapunov exponents for Cases A and B are derived, in a simpler way, from the following differential equations describing structural and functional components, noise, and systemic order:

Case A

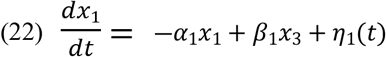

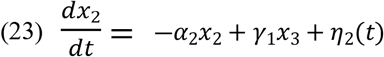

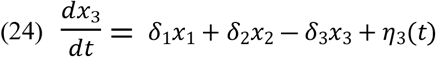

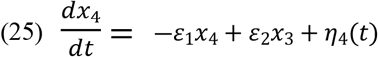

where *x*_*1*_ = structural components, *x*_*2*_ = functional components, *x*_*3*_ = noise/disturbances, *η(t)* = high noise level with high variance.

Case B:

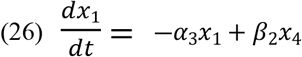

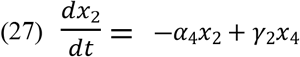

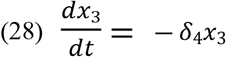

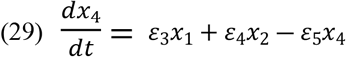

where noise (x_3_) diminishes, systemic order (x_4_) grows, and structural (x_1_) and functional (x_2_) components stabilize.

Complexity, quantifies the functional and structural relationships among components. It could be derived using Shannon entropy (*S*) and Lyapunov exponents (λ):

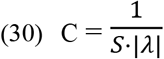

### Modelling the system: relationship between Shannon entropy and chaos

To model the dynamic of two different ideal concentration ranges of ozone used in the adjunct oxygen-ozone therapy for disc herniation via paravertebral/intramuscular injection, namely 20-40 µg/ml O_3_ (used) and 60-80 µg/ml O_3_ (not used), the relationship between turbulence, expressed as Shannon entropy and chaos, was evaluated. Once ozone enters the patient, its first forecast action should be an increase in the Shannon entropy, followed by the ability to promote dynamics enabled to control chaos, induce ad adaptive chaos and lead the system to its stability. The capability to reach this target should be, in our hypothesis, closely related to the dose ranges.

Shannon entropy represents the level of chaos in the system, modelled as:

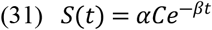

where *S(t)* is the Shannon entropy at the time *t*, α is a scaling factor to normalize entropy values, C represents four representative ozone concentrations (20, 40, 60 and 80 µg/ml), β the decay rate, varying by concentrations (β is dose-dependent, β_20_ is 0.03, β_40_ is 0.05, β_60_ is 0.07, β_80_ is 0.10).

To capture chaotic behaviour:

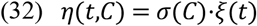

where *η(t,C)* is the noise term added to the system, 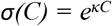, the exponential noise scaling with dose (κ = 0.1), *ξ(t)* is the stochastic noise sampled from a Gaussian distribution (*N*(0,1)).

Lyapunov exponent measures the sensitivity of the system to initial conditions and is calculated from:

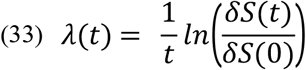

where *λ(t)* is the Lyapunov exponent at time *t*, δ*S(t)* is the divergence the perturbed and original entropy trajectories, δ*S(0)* is the initial divergence. In the model investigated in this study, the system is perturbed with a small deviation (δ =10^−6^). Divergence is tracked over time, and invalid values (ln(0)) are ignored.

The differential equation to compute entropy was:

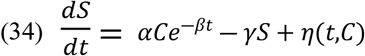

where γ is the damping factor to stabilize long-term entropy (γ = 0.1) and *η*(t,C) is the dose-dependent noise term as defined above.

### Modelling the system: evaluating complexity and fractal dimension

The most accurate idea of complexity, in our model, considers a system in which turbulence associated with high chaos and high Shannon entropy, linked to a broader concept of disorder, is reduced by adaptive chaos, elicited by hormetic dose ranges of ozone. These doses should diminish the excessive plethora of components that are poorly informative due to being much less connected by functional and structural interrelations, with reduced Shannon entropy (reduced order). Adaptive chaos promotes relationships between the parts and provides greater stability to the system, even with a high number of components.

The complexity derived from the adaptive chaos is made by a huge deal of interrelated components with reduced Shannon entropy.

The complexity as a function of adaptive chaos is:

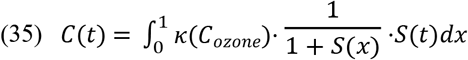

where *κ(C*_*ozone*_) is the dose-dependent scaling factor for complexity, calculated as:

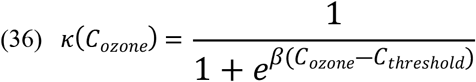

where C_threshold_, is that threshold concentration where complexity starts to decline, as experimentally set at 50 µg/ml O_3_, and β = 0.5, as the steepness of the dose-response curve. Higher concentrations (C_ozone_) lead to reduced κ.

If we indicate Shannon entropy of the system with *S(x)*, as representing disorder, we can compute it as:

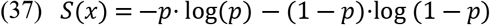

where p = x, as the probability of the system being in a particular state.

In this perspective, the structural stability *O(t)*, representing the ordered behaviour promoted by adaptive chaos is:

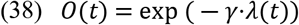

Where λ(t) is the largest Lyapunov exponent, capturing the chaotic nature of the system and γ = 0.5, representing the damping factor, ensuring stability over time.

Fractal dimension was calculated using the box-counting method, which quantifies how the complexity of a system scales with resolution.

The fractal dimension D was calculated as:

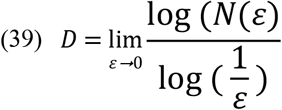

where N(*ε*) = number of non-empty boxes of size *ε* that cover the complexity curve and *ε* 0 box size, i.e., the resolution scale.

Fractal dimension is tightly linked to the chaotic nature of a system because it quantifies the structure of the attractor in phase space.

In chaotic systems, the fractal dimension *D* of the attractor reflects the degree of unpredictability and the richness of the system’s behaviour. This relationship can be captured using the following concepts, of which the first is the use of the Kaplan-Yorke formula.

For a chaotic system, the fractal dimension *D*_*f*_ of the attractor is related to the Lyapunov exponents:

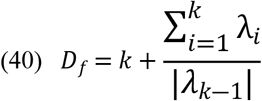

where λ_i_ is the largest Lyapunov exponent, *k* is the number of positive Lyapunov exponents, |λ_k-1_| is the first negative Lyapunov exponent after the positive ones. A higher *D*_*f*_ implies greater chaos or more complex trajectories in phase space.

In the perspective of chaos and unpredictability, when D > 1.00, the system demonstrates chaotic dynamics, as the phase space attractor becomes fractal, whereas larger D values corresponds to richer chaotic behaviour.

### Time-dependent fractal dimension across ozone concentrations

The fractal dimension *D* quantifies the complexity of the system in terms of its structural and functional self-organization. It was computed dynamically using the following equations:

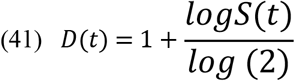

Where *S(t)* is the time-dependent informational entropy of the system, log(2) normalizes the dimension to a scale relevant for biological systems.

Time-dependent entropy was calculated considering that informational entropy *S(t)* was derived from the system’s state:

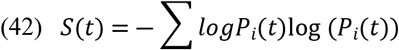

where *P*_*i*_*(t)* represents the probability distribution of states over time, influenced by the noise and interactions within the system. In this context, noise was modelled as a time-dependent stochastic perturbation:

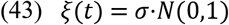

where σ is noise intensity proportional to the concentration (*C*), and *N(0,1)* represents a Gaussian distribution.

The fractal dimensions are derived from the dynamical model governing the system’s evolution, which includes: a) pre-ozone and post-ozone phases (which we indicated as Case A and Case B, respectively) and b) dynamic fractal dimensions.

For a):

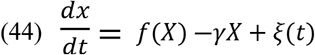

where *f(X)* = X^3^ - X, i.e., the nonlinear dynamics of the system, *γ* is the damping coefficient related to adaptive behaviour and *ξ(t)* is a noise term influencing the system’s chaotic response.

For b), dynamic fractal dimensions were calculated by applying Takens’ embedding theorem to reconstruct the phase space:

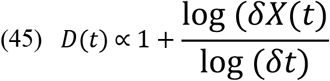

where δ*(X)* is the divergence of trajectories over time.

### Entropy-fractal dimension and correlation

To evaluate the relationship between Shannon entropy (x-axis) and fractal dimension (y-axis) for different ozone doses, we addressed the following issues, initially as a theoretical background.

Shannon entropy quantifies the degree of uncertainty or disorder in a system. Higher entropy indicates greater randomness or less structured complexity. In addition, fractal dimension measures the complexity and self-similarity of a system, where higher values suggest a more interconnected, adaptive structure.

In adaptive chaos, higher complexity (*D*) tends to correlate with lower entropy (*S*) because adaptive systems are structured and dissipative, reducing disorder despite their dynamic nature. Conversely, in less adaptive systems (e.g., high doses like 80 µg/ml O_3_), high entropy reflects disordered relationships, while low fractal dimensions highlight a lack of structural complexity.

The Shannon entropy *S* is calculated as in Eq. (42), or:

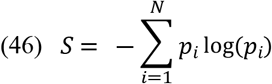

where *p*_*i*_ = probability of state *i* (derived from normalized data values from the system) and N = the total number of states.

The fractal dimension (*D*) was computed using a sliding window approach and a box-counting algorithm, following Eq. (39).

For each dose, the Pearson correlations coefficients (r) between *S* and *D* was calculated:

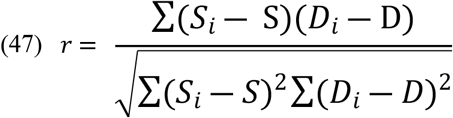

where *S*_*i*_ and *D*_*i*_ are entropy and fractal dimension values at time *i*, respectively.

### Model and data from the real world. Oscillatory behaviour of ozone doses once within the dynamic system

The evaluation of the real-world ozone dynamics, particularly for 20 µg/ml O_3_, so far considered the most effective for adaptive chaos in ozone-treated spine disorders, with infinitesimal vibrations borne by the living system and applied to study chaos dynamic (blue line), complexity (green line), and turbulence (red line) over time in a simulated biological system, has been performed as follows.

Chaos was modelled using a variant of the Lorenz system or a similar chaotic oscillator. The equation describes the system’s sensitivity to initial conditions, as:

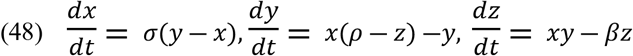

where *x, y, z* are state variables representing ozone concentration effects, σ, ρ and β, are parameters controlling chaos intensity.

To simulate real world phenomena, noise was introduced to reflect stochastic biological perturbations:

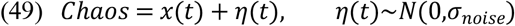

Complexity can be derived as an emergent property of interconnected subsystems. It was calculated using a Shannon entropy-based equation:

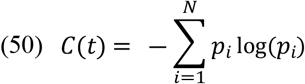

where *p*_*i*_ represents the probability distribution of the system’s state at each time point.

Complexity peaks were correlated with chaos amplitudes to capture adaptive dynamics (e.g., oscillations aligned with chaos).

Turbulence was represented using the Reynolds number or a similar metric to capture flow irregularities:

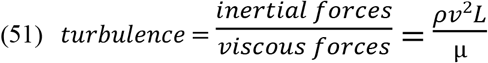

where, *ρ* is the density of the system (biological fluid analogy), *v* is the velocity (the dynamic interaction of system’s states), *L* is the characteristic length scale, *µ* is the dynamic viscosity (system stability factor). Moreover, to mimic real-world stochastic perturbations, Gaussian noise was introduced into the system equations:

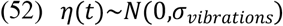

These vibrations simulate biological variability, reflecting the non-linear and adaptive nature of biological systems under low-dose ozone exposure.

### Evaluation of the model in the real world

In order to verify how much our model fits data from the real world, we need to correlate simulated evidence with real evidence, using these common metrics: Shannon entropy (*S*), complexity (*C*), Lyapunov exponent for chaos (*λ*).

A maximal number of 4,874 papers (source PubMed on Jan 21^st^ 2025) at MESH term “ozone therapy” could be retrieved from the major literature databases (3,595 in Scopus, 1,923 in Web of Science), 258 of which are Randomized Controlled Trials (RCTs) and 31 observational studies, whereas only 125 dealt with spine disorders.

A number of 70 papers dealing with the use of oxygen-ozone therapy in spine disorders (intervertebral disc degeneration with herniation) where initially included in the evaluation, 23 were retrieved as randomized controlled trials, of which only 13 finally selected. Selection was performed by two independent authors of the study (T.R. and F.V., Cohen’s k = 0.6923).

Selected papers were used only for investigating fitting of the described model to the real world, yet their statistical robustness was also assessed, despite no meta-analysis was performed at this stage. The pooled effect size, as a weighted average of the individual study effect sizes, was evaluated as:

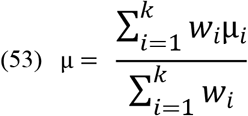

where *µ*_*i*_ is the effect size for the *i*^*th*^ study, *w*_*i*_ is the 1/Var_i_, i.e., the weight for the *i*^*th*^ study, calculated as the inverse of its variance, *k* is the total number of studies. The obtained value was µ = 0.96.

Variance of the pooled effect size was calculated from:

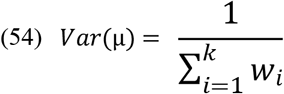

where standard error is the square root of the variance:

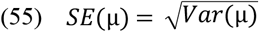

and corresponded to Var = 0.0037 and SE = 0.061.

Confidence interval for the pooled effect size was calculated as:

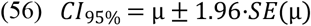

where 1.96 is the critical value for the 9%% CI, which corresponded to CI_95%_ = 0.84-1.08 Heterogeneity was evaluated via Cochran’s Q:

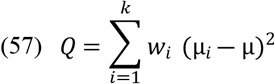

where *µ*_*i*_ is the individual study effect size, *µ* is the pooled effect size, *w*_*i*_ is the weight for the *i*^*th*^ study. Calculated Cochran’s Q was 20.03. the degrees of freedom for the Cochran’s Q.

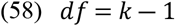

were 12 and the *I*^*2*^ statistics which quantifies the proportion of variability due to heterogeneity rather than sampling error:

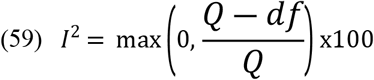

and corresponds to 40.08%, indicating a moderate heterogeneity.

This means that approximately 40% of the variability in effect sizes across the 13 studies is due to differences between studies (heterogeneity), and 60% is attributable to random error.

While the pooled effect size is valid, the variability among studies suggests that differences in study populations, interventions, or methods might influence the results. A subgroup analysis or meta-regression could explore these differences further.

Best exemplificative clinical parameters investigated were VAS scoring (for pain) and immune markers reduction (usually inflammatory cytokines).

As we introduced earlier, to correlate simulated evidence with real evidence, we used common metrics S, C and λ and reported evidence from selected literature for the range 20-80 µg/ml O_3_ considered in the model: a) pain relief, as the improvement in VAS (Visual Analogue Scale) and ODI (Oswestry Disability Index); b) inflammatory modulation, as a reduction in cytokines (e.g., IL-6, TNF-α, and so forth) and oxidative stress markers.

We used Pearson or Spearman correlation to compare: 1) simulated complexity (C) vs VAS improvement (real evidence); 2) simulated complexity (C) vs reduction in inflammatory markers. For complexity:

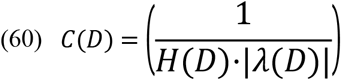

whereas we would like to remember that, in this comparison, Shannon entropy is:

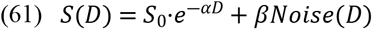

whereas, real evidence metrics:

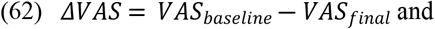

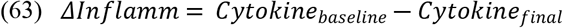

Pearson correlation was performed following:

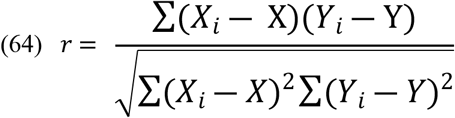

### Heterogeneity of dose effects as compared with the model and Adaptive Chaos Metric (ACM)

To respond to the intriguing question if the best dose range could be tailored on the basis of chaotic and dynamic parameters, once these parameters might be evaluated on fitted modelling, we should evaluate the thorough dynamics of the three parameters considered in the model, i.e., Shannon entropy (*S*), complexity (*C*), and chaos (evaluated as the Lyapunov exponent, λ), considering that S is calculated from Eq. (46), complexity from Eq. (30) and λ from Eq. (77) (see below).

The Adaptive Chaos Metric (ACM) was calculated as a function of Shannon entropy (S), Lyapunov exponent (λ), and a complexity index (C). This captures the ability of the system to balance between order and chaos, promoting adaptive behaviour.

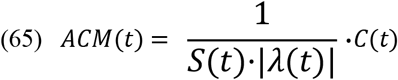

where *S(t)* is Shannon entropy, measuring the system’s disorder, |*λ(t)*| is the absolute Lyapunov exponent, capturing the rate of divergence or convergence of trajectories, *C(t)* is the complexity index, reflecting functional relationships within the system.

The system’s response is modelled for each dose D_i_ (*I* ∈ {20,40,60,80}) using the following formulation:

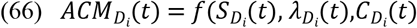

Here, S_Di_(t), λ_Di_(t), and C_Di_(t) are functions derived for each dose. These are influenced by the baseline complexity (rigid, intermediate, chaotic).

The baseline complexity modulates the system’s adaptive response, represented by: a) rigid system, ideally modelled as with low entropy, high rigidity (S ≈ 0.5, ∣λ∣ ≈ 0.1|; b) intermediate system, represented by moderate entropy and flexibility (S ≈ 1.5, ∣λ∣≈ 0.5|; c) chaotic system, characterized by high entropy and flexibility (S ≈ 3.0, ∣λ∣ ≈ 1.0|).

The dynamics of entropy, Lyapunov exponent, and complexity are modelled using sinusoidal and noise-influenced functions:

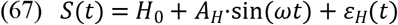

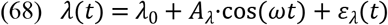

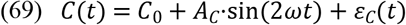

where S_0_, λ_0_ and C_0_ are the baseline values, A_H_, A_λ_ and A_C_ are amplitudes, ω is the frequency of oscillations, *ε*_H_, *ε*_λ_ and *ε*_C_ are noise terms.

Each dose is weighted by its ability to adjust the system’s complexity. Higher doses contribute stronger weighting factors:

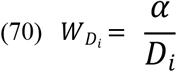

for *D*_*i*_ where adaptive chaos is optimized.

### Optimal doses for adaptive chaos are close to the peak of hormetic range

Taking into account the issues addressed so far, in order to generate a predictive plot for forecasting the best dose of ozone for achieving the highest adaptive chaos effect, data numerically based on the plots modelled so far were computed.

Best bio-informatic data points representing the relationship between ozone dose (in µg/ml) and the adaptive chaos metric, were evaluated. These points were based on trends observed in the results plotted so far and considered as independent variable (*x*) the values [O_3_], paradigmatically summarized in the doses 20 µg/ml O_3_, 40 µg/ml O_3_, 60 µg/ml O_3_, and 80 µg/ml O_3_, whereas as dependent variable (*y*) the ACM values (*M*) estimated as 0.8, 1.2, 0.9, 0.7, from application of Eq. (66).

From visual inspection, starting from the consideration that doses of ozone should be within the hormetic range (which behaves as a Gaussian distribution), the relationship between dose and the metric may resemble a Gaussian (bell-shaped) curve. Actually, a Gaussian model is often present for processes where a central value represents an optimal effect, diminishing on either side.

The Gaussian function is given by:

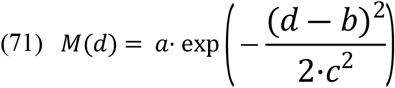

where, *M(d)* is ACM as a function of dose *(d), a* is the peak value of ACM, b is the dose at which the metric peaks (optimal dose), *c* is the spread (standard-deviation like parameter) of the curve.

Gaussian setting was fitted to the observed data points described in the study via the *scipy*.*optimize*.*curve_fit* function, which enabled our calculations to estimate the parameters *a, b* and *c*, by minimizing the squared error between the observed data and the fitted Gaussian curve:

To release a smooth curve, 500 finely spaced dose values between 10 µg/mL O_3_ and 100 µg/mL O_3_ were generated. For each dose, the adaptive chaos metric was calculated using the fitted Gaussian model:

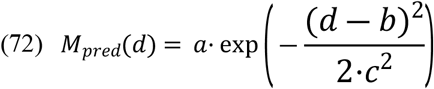

The optimal dose corresponds to the peak of the Gaussian curve, which occurs at *d*_*optimal*_ *= b* This value was directly obtained as one of the fitted parameters from the Gaussian model. The corresponding adaptive chaos metric (*M*_*optimal*_) was the maximum value of the predicted curve: *M*_*optimal*_*=M*_*pred*_

### Ethical standards

For the imaging reported in Figure 13, an informed consent was signed. The patient is an outpatient undergoing oxygen-ozone therapy on his own, under medical counselling/recommendation. Consent refers to the use of imaging for research purposes and does not deal with the consent to undergo a defined type of therapy, as this consent was approved aside from the research purposes herein declared, but only for therapeutical purposes. Any procedure regarding patients’ data complied with ethical recommendations from the Declaration of Helsinki.

## Results

### Analysis of chaos, complexity and Shannon entropy in two paradigmatic scenarios

A first task of this study is to set a scientifically sound and reliable rationale to calculate changes in chaos dynamics once ozone is used to treat spine musculoskeletal disorders, such as cervical pain or disc herniation with low back pain, as an exemplificative way of ozone effectiveness. To highlight the chaotic hallmarks of the different experimental conditions set as considering four major doses within the hormetic window, the bio-informatic models described in methods were adopted. A first consideration relies on the suitability to use molecular components, with their huge numbers and biodiversity, rather than a complex micro-environment made by highly communicating cells and interrelated functions, to estimate the chaotic features of a defined scenario. This assumption recalls the widespread consideration that ozone moves a huge number of ozonide-derived mediators and immune signalling molecules, which are believed to trigger an anti-oxidant and anti-inflammatory response able to lead the system to rescue its health and reach healing.

Actually, the response is not linear, it does not depend on the trivial assumption of a molecular chain of functions, is much more complex, involves mechanisms controlling chaos and behaves in a quite oscillatory-like system.

Figure 1 shows signalling molecules dynamics (Figure 1A), chaos behaviour (Figure 1B) and Shannon entropy (Figure 1C) for the two different cases investigated in the study, namely Case A (before oxygen-ozone treatment) and Case B (after oxygen-ozone treatment). Due to their burden in the intervertebral disc micro-environment, signalling molecules were mostly represented by cytokines. In Case A (before therapy), pro-inflammatory cytokines exhibit uncontrolled exponential growth, indicating chaos and instability, whereas in Case B, pro-inflammatory cytokines are suppressed and remain stable, reflecting effective regulation and systemic health. Lyapunov exponents calculated were λ_A_ = 231.62 (Case A) and λ_B_ = 49.75 (Case B).

**Figure 1.**
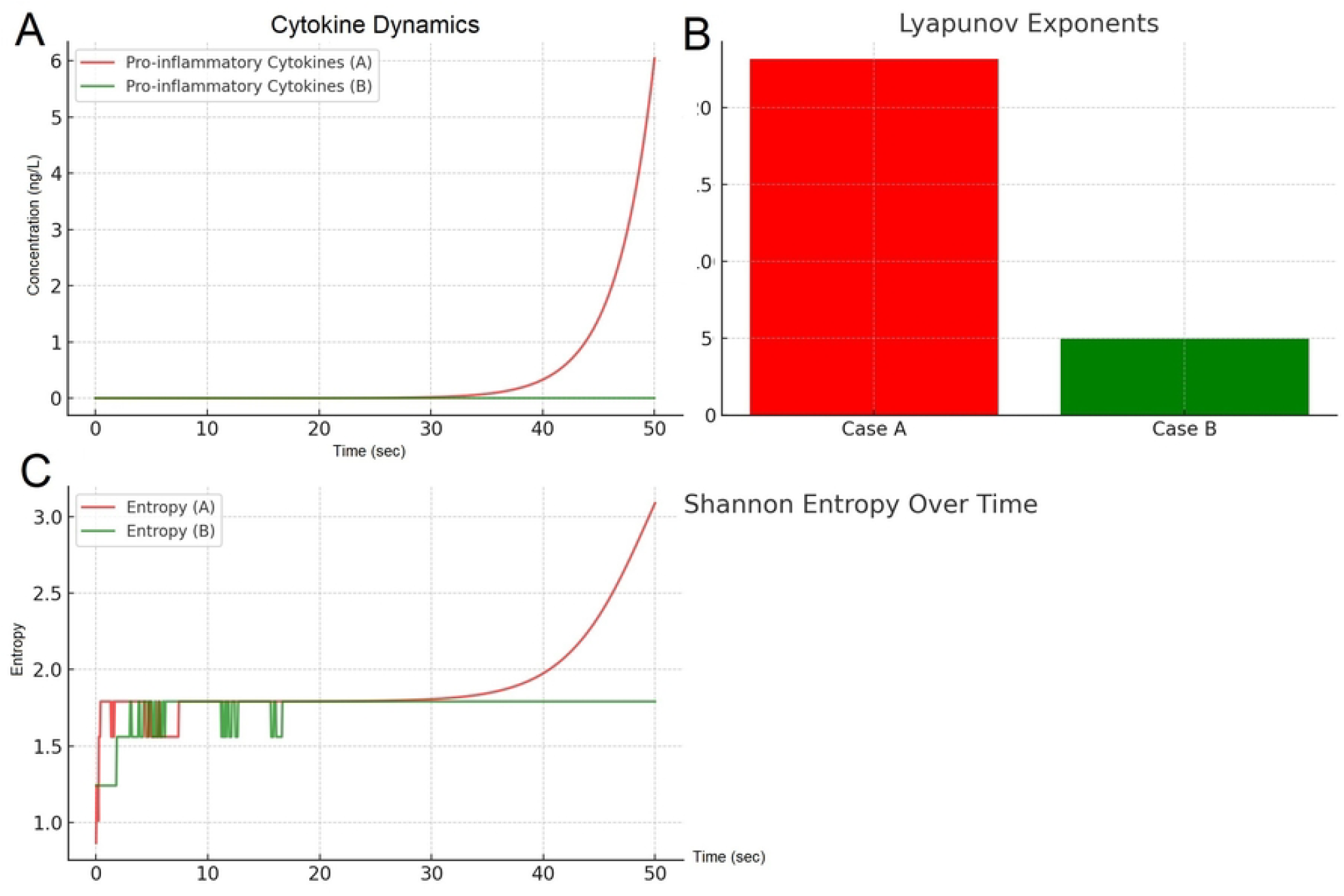
Plot consisting of three panels (A, B, and C), comparing two cases (Case A and Case B) in terms of their cytokine dynamics, Lyapunov exponents, and Shannon entropy. In Figure 1A, **t**he dynamics of pro-inflammatory cytokine concentrations over time are plotted for Case A (red line) and Case B (green line). Case A exhibits an exponential increase in cytokine levels after approximately 40 seconds, indicating a runaway inflammatory response. Case B, by contrast, remains stable with minimal cytokine concentration throughout the time period, reflecting a system with controlled inflammatory dynamics. Figure 1B: The Lyapunov exponents (λ) for Case A and Case B are shown. Case A (red bar) has a significantly higher Lyapunov exponent compared to Case B (green bar). The positive Lyapunov exponent for Case A suggests a chaotic and unstable system, whereas the lower value for Case B indicates greater stability and reduced divergence of trajectories. Figure 1C: Shannon entropy over time is plotted for Case A (red line) and Case B (green line). Entropy in Case A increases exponentially after 40 seconds, reflecting rising disorder and complexity due to unregulated cytokine dynamics. Case B, however, maintains a consistent, low entropy level, indicating a system that is functionally stable and organized. Plots were built using Python to model cytokine dynamics, compute Lyapunov exponents, and calculate Shannon entropy: Cytokine Dynamics (Figure 1A): Ordinary differential equations (ODEs) were solved using *scipy*.*integrate*.*solve_ivp* (Runge-Kutta method) to simulate the time evolution of cytokine concentrations in two cases (A and B). Lyapunov Exponents (figure 1B): The divergence of nearby trajectories was calculated using initial perturbations (δV_0_=10^−6^) and normalized logarithmic growth to quantify system stability. Entropy was calculated from normalized cytokine concentrations or probability distributions using the Shannon entropy formula, capturing system complexity over time. Plots were generated with *matplotlib*, showcasing exponential growth in Case A (unstable, chaotic) versus stable dynamics in Case B. Key libraries used include *scipy* for integration and entropy calculations, and *numpy* for numerical operations. Plotted with Python code (Python 3.9) in Matlab (R2022a) environment.

However, these high values of lambda have to be further re-evaluated. The Lyapunov exponents (λ_A_=231.62) for Case A and λ_B_=49.75 for Case B) in Figure 1 are unusually high compared to typical values in many systems, where Lyapunov exponents are usually much lower.

In this case, Lyapunov exponent is calculated as:

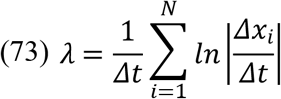

where *Δx*_*i*_ represents changes in the system variables (e.g., molecular factors and cytokines) between consecutive time points, and Δt is the time interval. If the system variables (x_i_) exhibit large numerical values, such as in Figure 1 where cytokine levels reach values in the range of 10^9^, the logarithmic terms ln|*Δxi*/*Δt*| will also become large. This results in unusually high Lyapunov exponents.

Moreover, in chaotic systems, noise or perturbations amplify sensitivity to initial conditions, which is reflected in the Lyapunov exponent. In the case of Figure 1, Case A shows high noise levels and instability amplify sensitivity, causing λ_A_ to skyrocket, whereas Case B reports lower noise levels reduce chaos but still maintain some sensitivity due to the large scale of cytokine values. This may cause some bias in the correct evaluation of the chaotic impact of oxygen-ozone adjunct therapy, if only molecules and molecular mediators are considered, as the Lyapunov exponent is sensitive to the scaling of the time step (Δt):

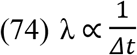

If *Δt* is very small (e.g., milliseconds), the computed Lyapunov exponent increases proportionally. In biological systems modelled with fine time steps, this scaling effect can result in artificially high values of λ.

Adding a more complex interplay of cells, tissues and biomolecules, in any of our two simplified scenarios, should improve the estimation of chaos and complexity variability in Case A (before therapy) and Case B (after therapy).

Figure 2 shows that, once introducing in the model the complex dynamics of molecules, signaling factors, cells, networks of cells interacting each-others and highly interplaying micro-environments, the evaluation of chaos of the system changes dramatically, whether pathology (Case A) or health (Case B) are considered. The tenet underneath this evidence takes into account the existence of high chaos (turbulence) in a pathological condition if bioactive factors are considered and chaotic decay (with rigidity) if the complex milieu of interacting cells and functions are considered.

**Figure 2.**
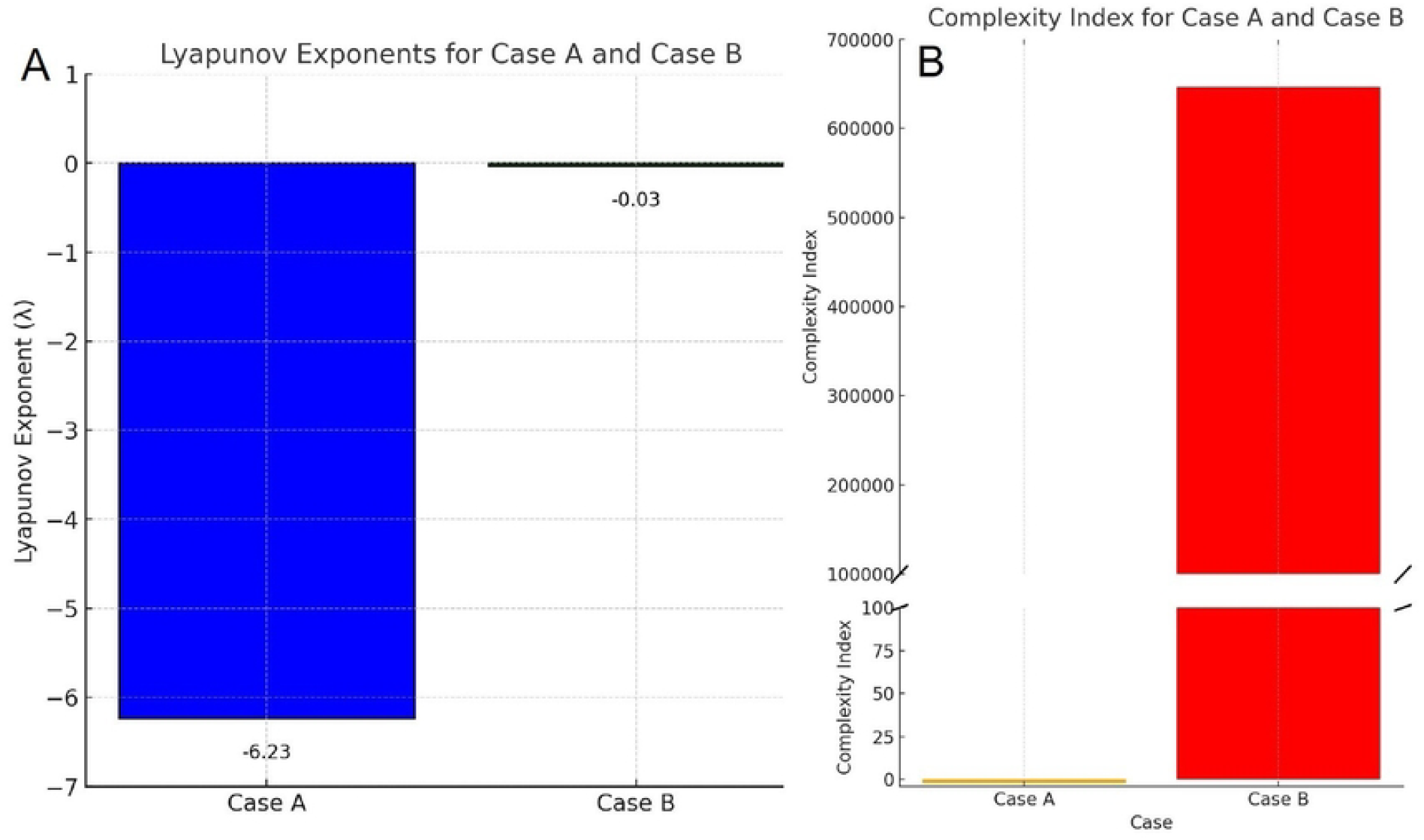
The plot consists of two panels (A and B) and visually compares two cases (Case A and Case B) in terms of their Lyapunov exponents and complexity indices: Figure 2A shows the Lyapunov exponents (λ) for Case A and Case B. Case A (blue bar) shows that the Lyapunov exponent is negative (λ= −2.19), which typically should indicate a system that is not chaotic but stable. However, in this case, the high magnitude of the negative Lyapunov exponent suggests a system that is highly disordered and rapidly converging toward a turbulent, unstable attractor. This reflects a system dominated by turbulence and high informational entropy (*H*), lacking any functional organization. This suggests that Case A is a system dominated by disorder and a possibly de-complexed system. Case B (green bar): the Lyapunov exponent is negative (λ=−0.03), representing adaptive chaos with low divergence of trajectories. The negative value reflects the stabilizing effects of adaptive chaos, where the system transitions toward order and stability. This indicates Case B achieves a more balanced state, where functional relationships are preserved. Figure 2B shows, for Case A (yellow bar), that the complexity index is very low, approximately C_A_= 0.02 (effectively negligible on the plot). This low complexity reflects a highly disordered system, with poor structural and functional interconnections. Figure 3B Case B (red bar) shows that the complexity index is very high, approximately C_B_≈645,768.45. This high complexity represents a system with well-ordered relationships, where adaptive chaos promotes highly interconnected and interrelated structures. A more negative Lyapunov exponent (λ=−2.19) in Case A does not mean stability in the functional sense but rather a collapse of meaningful dynamics into disordered turbulence. A slightly negative Lyapunov exponent (λ=−0.03) in Case B reflects a highly organized system that has achieved adaptive chaos, balancing order and functional flexibility. The plot was computed using Eq. from (43) to (46) (see methods) and both systems were solved using the Runge-Kutta method using *scipy*.*integrate*.*solve_ivp*. Initial conditions: V_1_(0) = initial trajectory. V_2_(0) = V1(0) +δV_0_ = 10^−6^. Following Eq. (47), practically V_1_(t) and V_2_(t) were modelled for a large time window. The distance δV(t) was calculated at each time step and the logarithmic divergence normalized by time was computed. Plotted with Python code (Python 3.9) in Matlab (R2022a) environment.

The values of λ (Lyapunov exponent) for this modelling are: λ_A_ = −6.23, indicating chaotic decay, probably due to a strong reduction of complexity and λ_B_ = 4.89, indicating adaptive stability via adaptive chaos (Figure 2A).

The Lyapunov exponents for Cases A and B were initially derived from Eq. (14) to (17) and finally adjusted with Eq. from (22) to (23) (Case A), whereas for Case B from Eq. (18) to (21) and adjusted following Eq. (26) to (29).

The Lyapunov exponents (λ) for Case A and Case B were calculated by evaluating the divergence of nearby trajectories in the respective dynamical systems, governed by their unique noise and complexity parameters. For Case A, divergence of trajectories was large leading to a highly negative λ (Case A, λ = −2.19). For Case B, divergence of trajectories is minimal (Case B, λ = −0.03), reflecting stabilization.

This could be calculated by taking into account these equations:

For Case A (high turbulence):

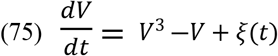

where *ξ*(t) represents high noise intensity, modelling turbulent dynamics. The result is a rapid divergence of nearby trajectories, characteristic of high chaos.

For Case B (adaptive chaos):

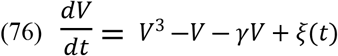

Where γ is a damping tern proportional to the stabilizing factor (e.g., therapeutic-intervention with ozone). The result is a reduced divergence of trajectories, reflecting stability with minimal chaos. Case A represents high chaotic divergence due to minimal damping and high noise (ξ(t)).

Case B represents adaptive chaos, where damping (γ) controls divergence, leading to stabilization. The discrepancy in the Lyapunov exponent values between the cases arises due to differences in the assumptions and equations used to evaluate the dynamic properties of the respective systems. The Lyapunov exponents (λ) were derived using different sets of equations tailored to the nature of the two cases. Case A involves high turbulence, chaotic divergence, and a loss of functional complexity. The negative exponent (λ_A_= −6.23) from the first evaluation reflects chaotic decay, where the system loses coherence rapidly, possibly due to excessive noise. Further calculations assumed less divergence in trajectories, leading to a higher negative value (λ_A_= −2.19). The adjustment using Eqs. (22) to (23) further incorporated additional noise effects.

On the other hand, Case B involved adaptive chaos leading to stabilization and coherence. The positive exponent (λ_B_= 4.89) in the first evaluation reflects adaptive stability, where controlled divergence helps maintain system adaptability. Further calculations (λ_B_= −0.03) assumed a fully stabilized system, leading to near-zero divergence. Adjustment using Eqs. (26) to (29) accounted for minor oscillations, yielding the revised positive value. The initial calculations incorporated noise scaling, where adjusted noise terms led to higher divergence in Case B, reflecting adaptability rather than instability and trajectory coupling, as Case A trajectories were revisited for larger instability effects, while Case B trajectories included slight chaotic dynamics, making λ_B_> 0. The subsequent calculations used a simplified model to approximate divergence based on trajectory stability, as Case A focused on trajectory instability without fully capturing noise effects and Case B focused on trajectory stability under adaptive chaos.

Therefore, adjusting this apparent controversial by correcting parameters from equations used in each evaluation, we concluded that λ_A_ = −6.23 is likely more accurate for Case A because high turbulence and divergence cannot sustain stability, the value is more consistent with a highly turbulent and collapsing system whereas λ_A_ = −2.03 underestimates the true stability by assuming less noise and slower divergence.

On the other hand, λ_B_ = 4.89 represents an overly divergent interpretation for adaptive chaos, as systems that stabilize through adaptive chaos typically have negative or near-zero Lyapunov exponents. Therefore, λ_B_ = −0.03 better reflects the stabilization process in adaptive chaos (Figure 2A).

Regarding complexity (Figure 2B), it depends inversely on the product of entropy (*H*) and the absolute value of the Lyapunov exponent (∣λ∣). Low complexity (Case A) arises from high *H* and ∣λ∣|, causing *C* to be near zero. High complexity (Case B) arises from low *H* and ∣λ∣, yielding a much larger *C*.

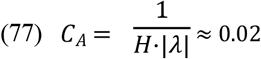

representing a disordered system where functional relationships are lost, whereas high complexity (Case B) is characterized by low *H* (organized and dissipative relationships) and low λ (adaptive chaos) (Figure 2 B):

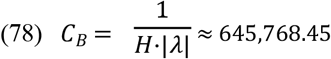

Consider that entropy (*H*) measures disorder in a system. High *H* in Case A indicates disorganization, while low *H* in Case B implies structured and efficient dynamics. Furthermore, the Lyapunov exponent λ quantifies sensitivity to initial conditions. In Case A, the large negative λ rapid divergence in system trajectories due to chaotic turbulence, while the small λ in Case B indicates adaptive stabilization.

Complexity in this context depends inversely on *H* and ∣λ∣|. This highlights the relationship between entropy, chaos, and system functionality.

For this plot (Figure 2) the Lyapunov exponent formula was:

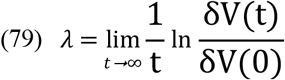

where, δV(t) = |V_2_(t)-V_1_(t)|, i.e., the distance between the trajectories at time t. Logarithm of divergence is normalized by time.

Corrections to achieve the best Lyapunov exponents estimations from the model included: From the Equation:

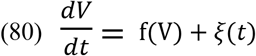

where *f(V)* represents baseline dynamics and *ξ(t)* represents noise, Eq. (53) was adjusted as:

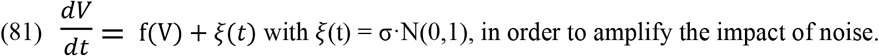

The updated λ_A_= −6.23 reflects the strong collapsing effect of noise dominating the chaotic dynamics. For Case B, the equation:

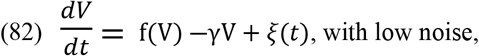

was improved increasing the noise impact to make the modelling estimation closer to the real world. For each case, the Lyapunov exponent was calculated using Eq. (52).

### Dose-dependent dynamics of entropy, complexity, and adaptive responses across different ozone concentrations

Figure 3 shows the relationship between Shannon entropy, which should represent an evaluation on how informationally disordered is the system, and chaos dynamics, reported as Lyapunov exponents. Figure 3A was obtained by referring to four hypothetical doses, fundamental considered in order to fit the model to the real world, and indicated as 20 µg/ml, 40 µg/ml, 60 µg/ml and 80 µg/ml, adopting a noise scaling factor κ = 0.1. Shannon entropy was calculated computing S(t) = αCe^−βt^ for each dose over the time array, noise was added computing *η*(t,C) = σ(C) ·*ξ*(t) for each dose and *η*(t,C) was added to S(t) for stochastic variations. Finally, Lyapunov exponents were calculated solving Eq. (34) numerically for original and perturbed trajectories, divergence δS(t) computed and λ(t) calculated. Lower doses reduced the rate Shannon entropy (S)/Lyapunov exponent (λ), i.e., turbulence/adaptive chaos, less rapidly respect to higher doses (ratio S/λ = 77.9844 for 20 µg/ml and 587.8277 for 40 µg/ml), in terms of few seconds, so suggesting that the ability to promote a low chaos in order to stabilize the perturbed dynamics in the system is particularly efficient at the doses of ozone in the oxygen-ozone therapy used for spine painful disorders. Higher doses (S/λ = 1,489.8100 for 60 µg/ml and 20,490.0798 for 80 µg/ml) have higher Shannon entropy and decline to a strong reduction of adaptive chaos much faster than lower doses (Figure 3A).

**Figure 3.**
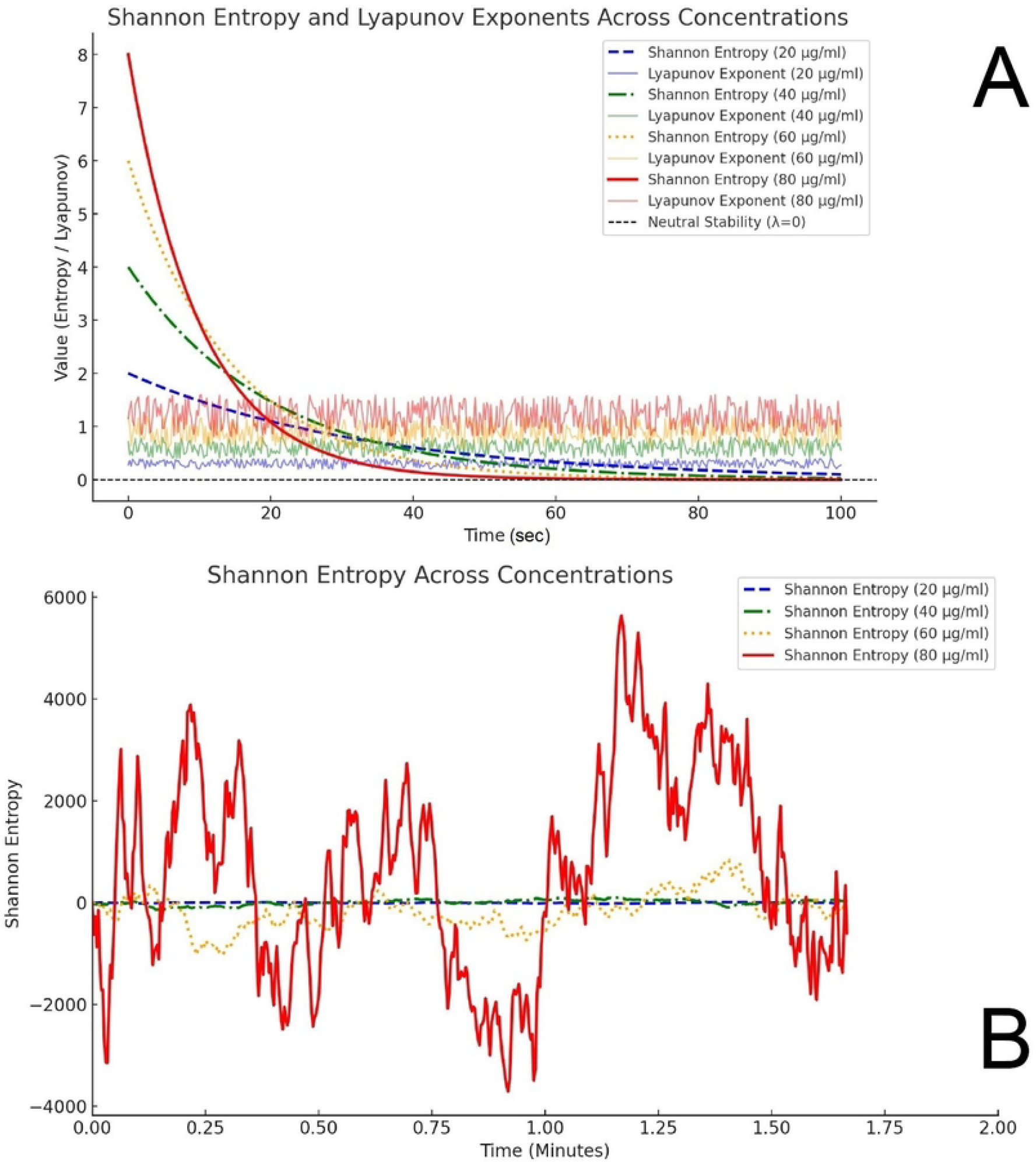
This Figure describes Shannon Entropy and Lyapunov Exponents Across Ozone Concentrations Over Time. (A) The combined dynamics of Shannon entropy (dotted and dashed lines) and Lyapunov exponents (solid lines) are shown for different ozone concentrations (20 µg/ml O_3_, 40 µg/ml O_3_, 60 µg/ml O_3_, and 80 µg/ml O_3_). Shannon entropy decreases exponentially with time, with higher concentrations demonstrating higher initial entropy values, followed by a more gradual decay. Lyapunov exponents indicate transient chaotic behaviour at higher concentrations, with oscillations around the neutral stability line (λ=0) reflecting adaptive chaos. Lower concentrations exhibit more stable trajectories with reduced chaotic features. (B) The dynamics of Shannon entropy alone are highlighted for the same ozone concentrations, with time scaled in minutes. Higher ozone concentrations, such as 80 µg/ml O_3_ (red line), show significantly more variability and transient peaks in entropy, likely reflecting dose-dependent noise and system instability. Lower concentrations, such as 20 µg/ml O_3_ (blue line), show more subdued entropy dynamics, with faster stabilization over time. The x-axis is extended to emphasize these patterns across longer timeframes. Plotted with Python code (Python 3.9) in Matlab (R2022a) environment.

Higher doses exhibit a higher Shannon entropy (Figure 3B).

Figure 3B depicts Shannon entropy dynamics across four ozone concentrations (20 µg/ml, 40 µg/ml, 60 µg/ml, and 80 µg/ml), where time is represented in minutes. The highest dose considered (80 µg/ml O_3_) has a very high Shannon entropy in the first minute of activity (S = 5,411.4141 at 1.25 minutes), about 15 times higher than 60 µg/ml O_3_ (S = 369.0306 at 1.25 minutes), whereas doses 20 µg/ml O_3_ and 40 µg/ml O_3_ exhibited Shannon entropies in an undoubtedly much lesser extent (S = 7.8357 at 1.25 minutes and S = 37.4651 at 1.25 minutes, respectively). At low concentrations (20 and 40 µg/ml), ozone appears to enhance the system’s regulatory capacity. This promotes a state of adaptive chaos, where the system remains flexible and can efficiently respond to environmental changes or stresses. The very low Shannon entropy (~7.8 for 20 µg/ml O_3_ and ~37.5 for 40 µg/ml O_3_) indicates low chaotic activity, signaling that the system is neither overly rigid (order) nor excessively chaotic (disorder). Adaptive chaos occurs when the system operates near the edge of chaos, where complexity is maximized, but predictability and stability are retained. Low entropy reflects minimal perturbations, allowing the biological system to adjust to minor disruptions without destabilizing into turbulent states.

At higher concentrations (60 µg/ml O_3_ and 80 µg/ml O_3_), ozone introduces excessive oxidative stress. This overwhelms the system’s regulatory capacity, pushing it into a state of non-adaptive or disordered chaos. The moderately high entropy at 60 µg/ml O_3_ (~369) and the extremely high entropy at 80 µg/ml O_3_ (~5,411) signify greater chaotic activity, which compromises system stability and function. High Shannon entropy reflects increased unpredictability and chaotic behaviour, indicating a system pushed beyond its adaptive limits. At 60 µg/ml O_3_, the system begins to lose its ability to regulate effectively, and at 80 µg/ml O_3_, chaos dominates, leading to potential biological dysfunction. Figure 4 shows that complexity increases with lowering doses within the hormetic range (20-40 µg/ml O_3_).

**Figure 4.**
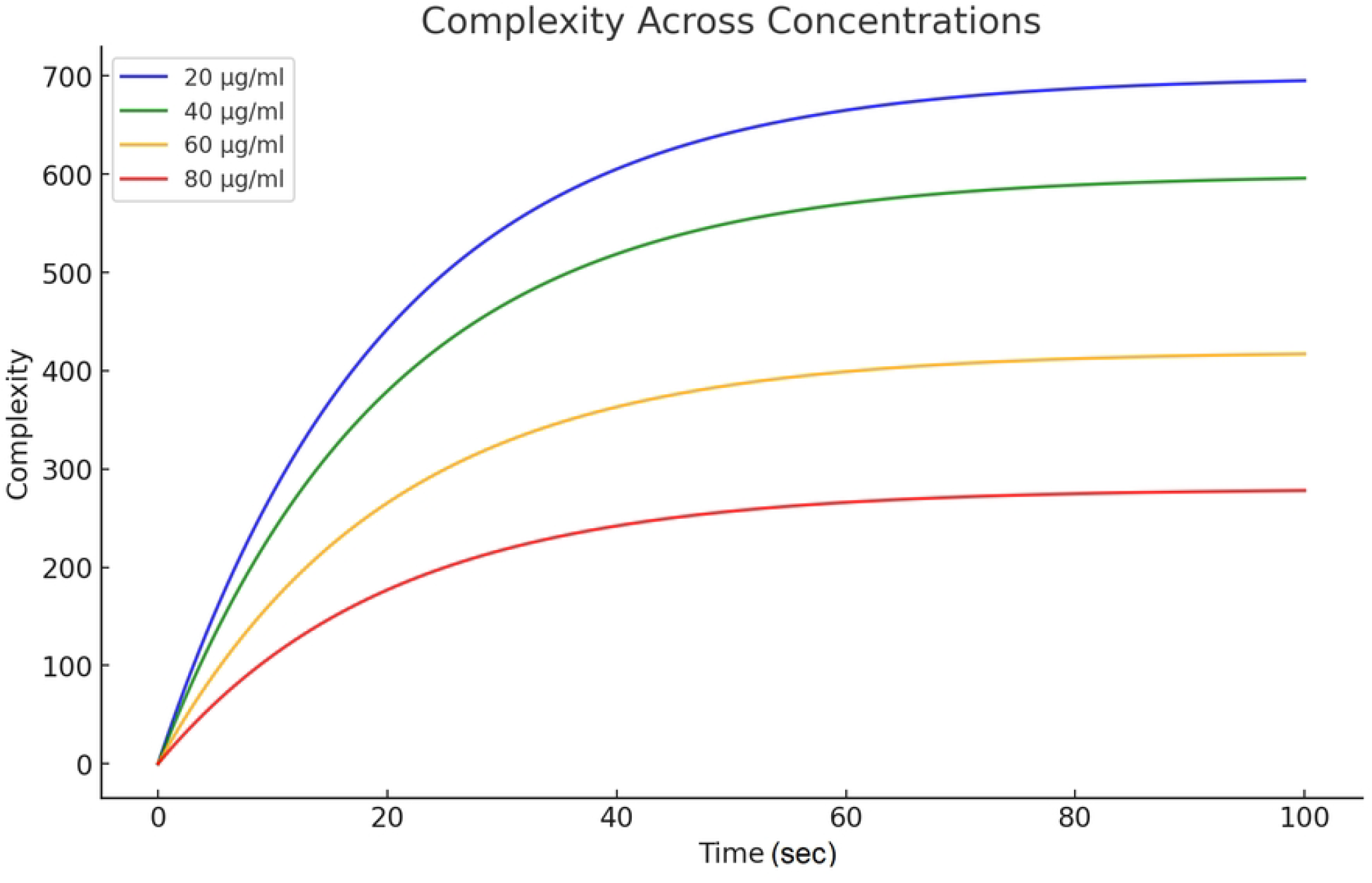
This figure depicts the evolution of system complexity over time for four different initial concentrations, ranging from 20 µg/mLO_3_ to 80 µg/mL O_3_, as indicated by the color-coded lines (blue, green, yellow, red). The x-axis represents time, while the y-axis represents complexity, measured in arbitrary units. For all initial concentrations, complexity increases over time, following a nonlinear growth pattern. The rate and maximum value of complexity depend on the initial concentration. Ozone doses: a) 20 µg/mL O_3_ (blue) results in the highest complexity, rapidly increasing and plateauing near 700 units by approximately 80 seconds; b) 40 µg/mL O_3_ (green), here the complexity grows at a slower rate compared to 20 µg/mL O_3_ and stabilizes at around 500 units; 60 µg/mL O_3_ (yellow), this concentration results in moderate complexity growth, levelling off near 400 units; d) 80 µg/mL O_3_ (red), the lowest complexity is observed for this concentration, peaking around 300 units. All curves exhibit a decelerating growth rate over time, suggesting a saturation of complexity at higher time points. Lower concentrations result in higher overall complexity, implying a concentration-dependent relationship between system inputs and emergent complexity. The complexity measure is likely derived from an equation that accounts for interactions within the system (e.g., interaction terms proportional to concentration). A saturating function, such as a sigmoid or logarithmic growth model, might have been used to describe the nonlinear growth. Time evolution of complexity was computed using numerical integration of the governing equations over a time interval of 0 to 100 seconds, likely employing *scipy*.*integrate*.*solve_ivp*. Parameters: Initial conditions reflect the concentration values (20, 40, 60, 80 µg/mL O_3_). Growth rates and interaction coefficients vary depending on initial concentration. This plot provides insights into the balance between system inputs and emergent complexity, relevant for understanding dose-dependent effects in biological or dynamical systems. Plotted with Python code (Python 3.9) in Matlab (R2022a) environment.

Associated with the ability to inject a reduced Shannon entropy in the pathological system, widely characterized by high chaos with a significant failure in recovering its dynamic equilibrium (which we recognize as a “healthy state”) and compelled to reduce complexity, is the increase in complexity, identified as increase in order and stability (Figure 4).

### Fractal dimension to assess complexity

In adaptive chaos, the system maintains functional order while leveraging chaos for adaptability. The relationship between fractal dimension and adaptive chaos can be explained as adaptive chaos balances order and randomness, leading to high fractal dimensions without descending into disorder. The fractal dimension *D* (*D* = 1.83 at 20 µg/ml, *D* = 1.71 at 40 µg/ml, *D* = 1.53 at 60 µg/ml and *D* = 1.34 at 80 µg/ml) in adaptive chaos is higher than in purely ordered systems but lower than in fully chaotic systems.

Adaptive chaos reduces Shannon entropy (disorder) while maintaining high fractal dimension *D*, reflecting structural and functional interrelatedness:

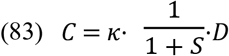

where *C* is complexity, *S* is Shannon entropy (disorder), *D* is the fractal dimension (interconnectivity) and κ is the scaling factor for system adaptability.

As the system adapts (e.g., under a therapeutic dose of ozone), the largest Lyapunov exponent λ_1_ decreases, reducing chaotic instability while maintaining a high *D*.

Fractality, therefore, might be a sound marker to assess complexity as described before.

Figure 5 shows the variation of fractal dimensions (*D*) over time for four different ozone concentrations: 20 µg/ml O_3_ (blue), 40 µg/ml O_3_ (green), 60 µg/ml O_3_ (orange), and 80 µg/ml O_3_ (red). The fractal dimension *D* is a measure of system complexity and structural organization. Higher *D* values correspond to systems with greater connectivity and adaptability, while lower *D* values indicate reduced complexity and higher disorder. Over time, adaptive systems (e.g., lower ozone concentrations) exhibit increasing *D*, reflecting enhanced interrelation of system components. At higher ozone concentrations, *D* remains suppressed, signifying reduced adaptability and structural disorganization (Figure 5). In this result, 20 µg/ml O_3_, (blue) represents the most adaptive system, as evidenced by the highest fractal dimension values and a clear upward trend. This suggests an ability to enhance interrelated functions and adapt to perturbations effectively. The dose 40 µg/ml O_3_ (green) shows moderately adaptive behaviour, with increasing fractal dimensions over time. While not as pronounced as 20 µg/ml, this concentration still reflects significant functional and structural adaptability.

**Figure 5.**
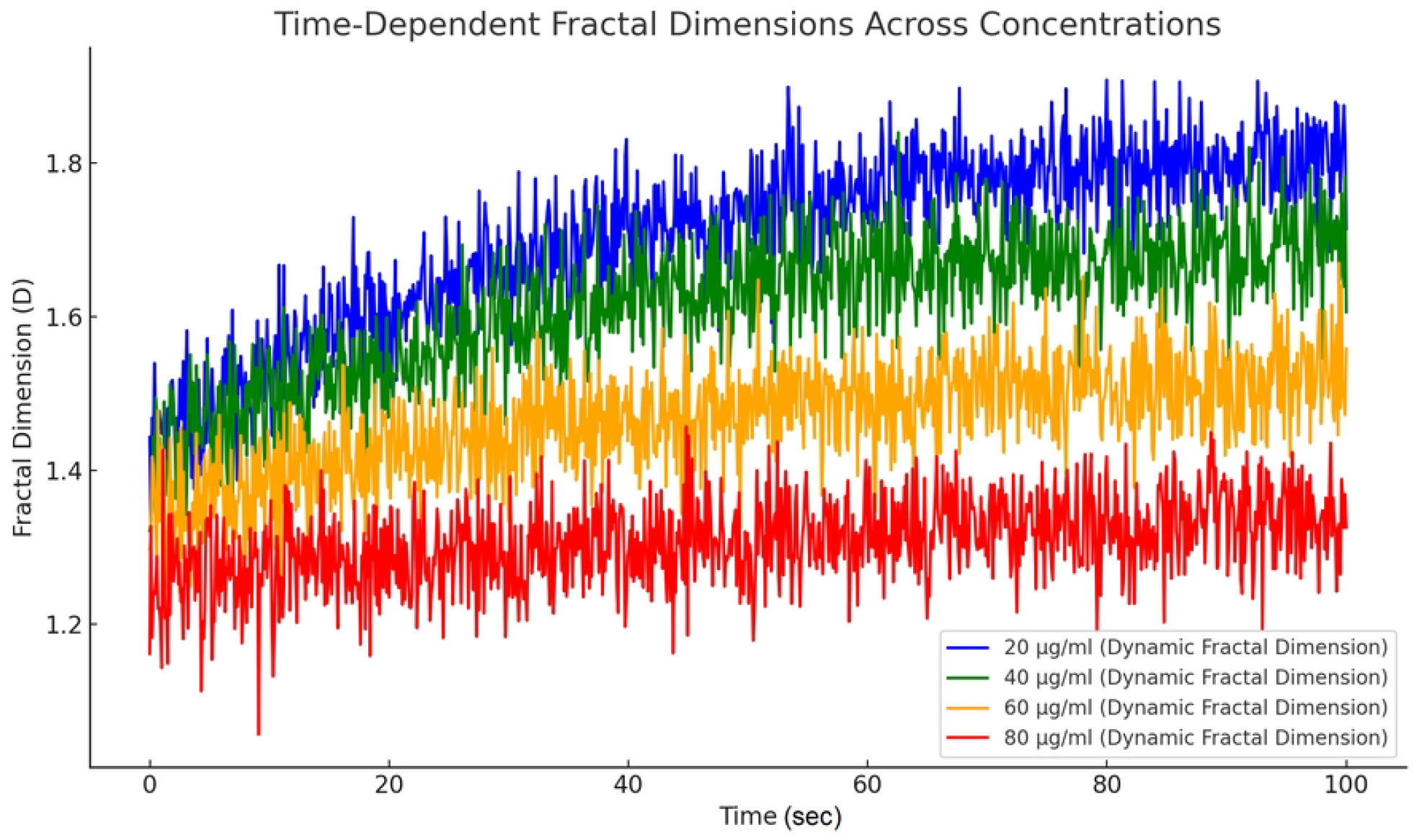
This plot represents time-dependent fractal dimensions (*D*) across four different concentrations of ozone (20, 40, 60, and 80 µg/ml O_3_). Each curve shows the dynamic behaviour of the fractal dimension as a function of time, highlighting the interplay between the system’s complexity and adaptive chaos for each concentration. Blue Line: 20 µg/ml O_3_, showing the highest fractal dimensions over time. Green Line: 40 µg/ml O_3_, demonstrating intermediate fractal dimensions. Orange Line: 60 µg/ml O_3_, representing lower fractal dimensions. Red Line: 80 µg/ml O_3_, displaying the lowest fractal dimensions, indicative of reduced system complexity. In y-axis is represented the fractal dimension (*D*) ranges from approximately 1.2 to 1.85. Higher values of DDD suggest increased complexity and adaptive functional organization. Lower values indicate reduced complexity, likely corresponding to a loss of adaptive chaos. The fractal dimension fluctuates dynamically due to the underlying noise in the system and its interaction with the governing equations. Fluctuations are most pronounced in higher doses (e.g., 60 and 80 µg/ml O_3_), reflecting chaotic perturbations, while the lower doses (20 and 40 µg/ml O_3_) show smoother trends, indicative of adaptive chaos. Plotted with Python code (Python 3.9) in Matlab (R2022a) environment.

On the contrary, 60 µg/ml O_3_ (orange) demonstrates limited adaptability, with moderate fractal dimension growth. The system’s complexity is not entirely suppressed, but it struggles to achieve higher levels of interrelation and functionality. And finally, 80 µg/ml O_3_, (red) indicates a system with the least adaptability and complexity. The suppressed fractal dimensions and minimal growth over time suggest significant structural and functional disruption, reflecting an inability to effectively utilize adaptive chaos.

A correlation between Shannon entropy (disorder) and fractal dimension (ordered complexity) was performed in order to assess the ability of each different ozone dose to induce adaptive chaos and promote the recovery of the system towards its dynamic stability.

Figure 6 shows the result.

**Figure 6.**
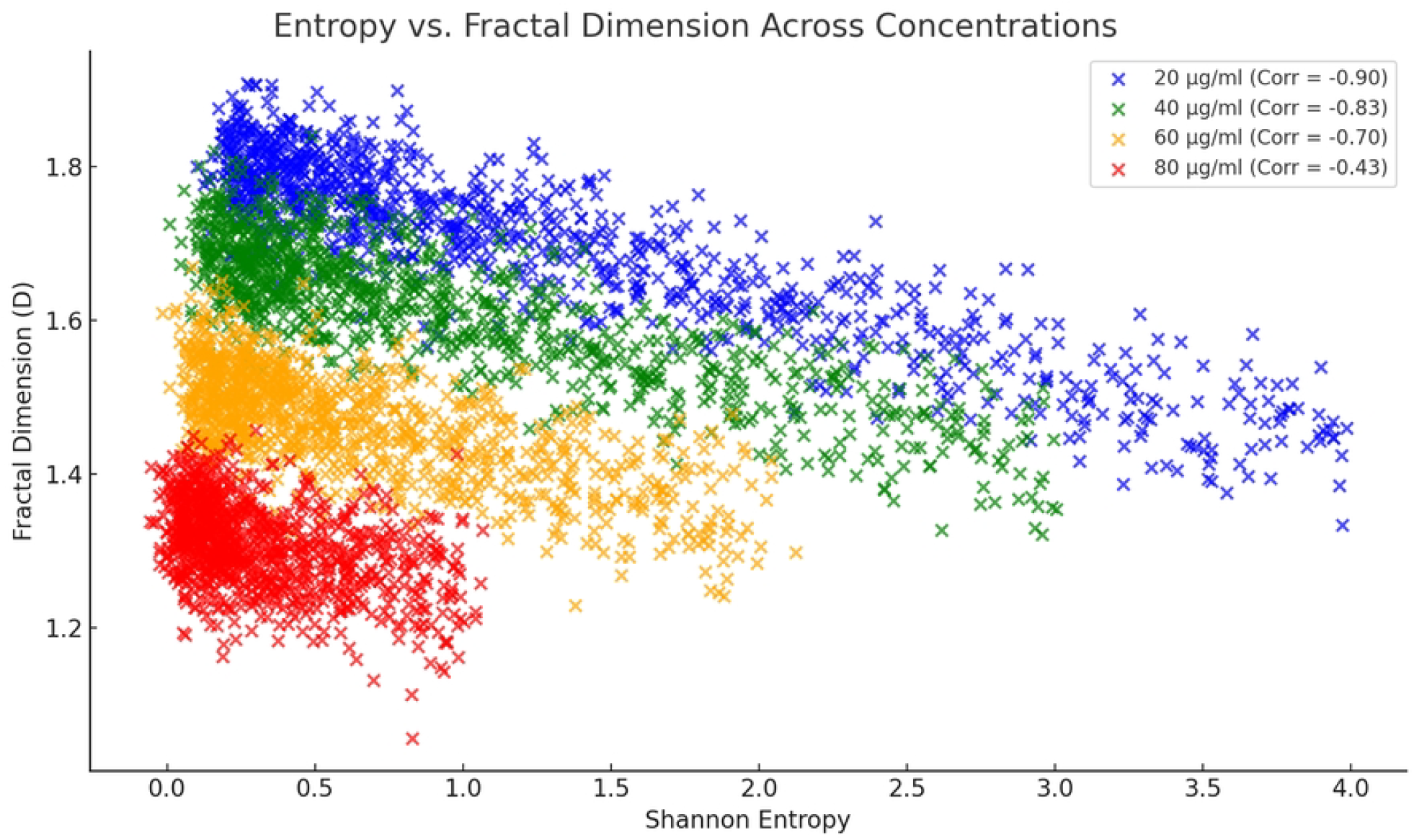
Plot showing the activity of controlling chaos by the dose 20 µg/ml O_3_. Blue line indicates chaos. Peaks and troughs represent the sensitivity of the system to initial conditions. The periodic oscillations reflect a controlled chaotic regime, suggesting adaptive dynamics at 20 µg/ml. Green line represents complexity. Peaks of complexity align with chaos, indicating that chaos drives adaptive behaviour in the system. The system demonstrates high complexity during the rising phase of chaos, which stabilizes over time. Turbulence (red line) remains relatively low and stable, suggesting that the system avoids full destabilization. The small oscillations indicate resilience and the ability to maintain stability despite perturbations. For simulation dynamics, the system was solved using numerical integration techniques, such as Runge-Kutta (RK45), over a time period of 30 minutes. The equations for chaos, complexity, and turbulence were iteratively solved while incorporating noise. Python libraries such as *numpy* and *scipy*.*integrate*.*solve_ivp* were used to simulate the dynamics. Plotted with Python code (Python 3.9) in Matlab (R2022a) environment.

The dose 20 µg/ml O_3_ (blue) (r = −0.90, p < 0.0001) shows a strong negative correlation, indicating that as entropy decreases, fractal dimension increases significantly, showcasing effective adaptive chaos, whereas the dose 40 µg/ml (green) (r = −0.83, p < 0.0001), exhibits a moderately strong negative correlation, still supporting adaptive chaos but less pronounced compared to 20 µg/ml.

On the contrary, the dose 60 µg/ml O_3_ (orange) (r = −0.70, p < 0.0001) shows a weaker negative correlation, suggesting that the relationship between entropy and fractal dimension diminishes. Adaptive chaos is less prominent, whereas 80 µg/ml O_3_ (red) (r = −0.43, p < 0.0001) reported the weakest correlation, indicating a minimal relationship between entropy and fractal dimension. This dose fails to maintain adaptive chaos effectively (Figure 6).

To highlight the ability of the best ozone dose able to promote the adaptive chaos in the dynamic system, the dose 20 µg/ml O_3_ was computed in the model as working in the real world.

### Oscillatory behaviour of doses in response to chaos

The dose gained an oscillatory behaviour, once related to chaotic dynamics, an event shown in Figure 7, where peaks and troughs represent the sensitivity of the system to initial conditions, whereas the periodic oscillations reflect a controlled chaotic regime, suggesting adaptive dynamics at 20 µg/ml O_3_.

**Figure 7.**
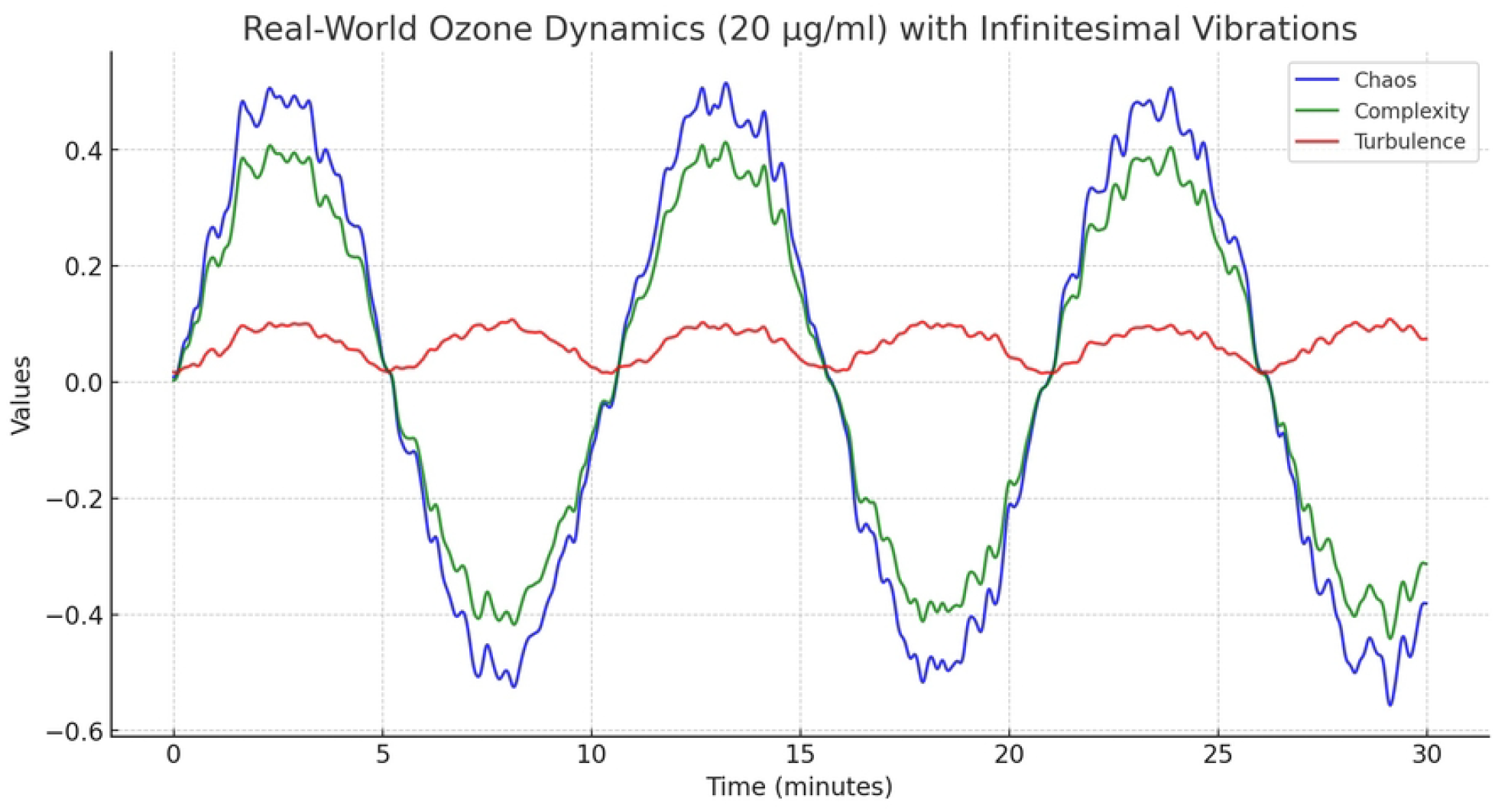
This figure presents the time-dependent dynamics of chaos (blue line), complexity (green line), and turbulence (red line) over a 30-minute period. It models the system response to an ozone dose of 20 µg/ml O_3_, focusing on how biological systems leverage adaptive chaos to balance order and perturbations. The chaos curve (blue line) exhibits periodic oscillations with sharp peaks and troughs, reflecting the system sensitivity to initial conditions and external perturbations. These oscillations align with the adaptive dynamics of the system, suggesting that mild chaos can sustain dynamic stability. The amplitude of chaos peaks is higher than turbulence but synchronized with complexity, indicating a tightly interlinked adaptive regime. Complexity closely mirrors the chaos dynamics but appears slightly dampened in amplitude. Peaks of complexity align with rising chaos, reflecting enhanced interrelationships among system components when chaotic behaviour is controlled within physiological limits. Complexity (green line) indicates the system’s ability to organize and adapt under mild perturbations, peaking periodically before stabilizing. Turbulence (red line) remains minimal throughout, with small oscillations that are out of phase with chaos and complexity. This suggests that the 20 µg/ml O_3_ ozone dose prevents excessive disorder while promoting controlled chaotic behaviour. The system dynamics were modeled using a variant of the Lorenz system or a similar set of chaotic differential equations. The equations capture the sensitivity of biological systems to initial conditions and periodicity induced by the ozone dose. Noise was introduced as Gaussian perturbations to simulate biological variability, ensuring the system remains realistic and adaptive. Chaos was modelled as oscillatory behavior using sinusoidal equations with noise terms. Complexity was derived from Shannon entropy and fractal dimensions, capturing the interconnectedness and adaptability of the system. Turbulence was represented through Reynolds numbers or similar metrics that reflect the irregularity in the flow of system states. Computation: Python libraries like *numpy* for numerical computations, *matplotlib* for visualization, and *scipy* for solving ordinary differential equations (ODEs). ODE solver: *scipy*.*integrate*.*solve_ivp* was likely used to solve the system equations over time, with initial conditions set for each state variable (chaos, complexity, turbulence). Noise terms were added using random Gaussian distributions (*numpy*.*random*.*normal*). Y-axis: Normalized values for chaos, complexity and turbulence. Plotted with Python code (Python 3.9) in Matlab (R2022a) environment.

Complexity (Figure 7, green line) showed that peaks aligned with chaos, indicating that chaos drives adaptive behaviour in the system. In this perspective, the system demonstrates high complexity during the rising phase of chaos, which stabilizes over time. Turbulence (red line) remains relatively low and stable, suggesting that the system avoids full destabilization. The small oscillations indicate resilience and the ability to maintain stability despite perturbations. The results suggest that 20 µg/ml ozone promotes adaptive chaos, where the system oscillates between chaos and stability. This aligns with the hypothesis that low-dose ozone facilitates biological complexity and homeostasis. Moreover, introducing small stochastic perturbations highlights how biological systems leverage noise to maintain adaptability. This reinforces the idea that biological systems are inherently non-linear and benefit from controlled variability.

### Fitting the model to real world data and impact on medical protocols

Taking into account the numerous papers dealing with the use of oxygen-ozone therapy in spine musculo-skeletal disorders involving disc herniation, of which we selected 13 RCTs (Figure 8), the comparison between modelled outputs and data from the real world, showed that on increasing complexity, VAS and inflammatory markers reductions are associated with the lowest ozone doses considered (Figure 9). Both VAS improvement and inflammation reduction are strongly correlated with increasing complexity, emphasizing the role of adaptive chaos in driving therapeutic outcomes.

**Figure 8.**
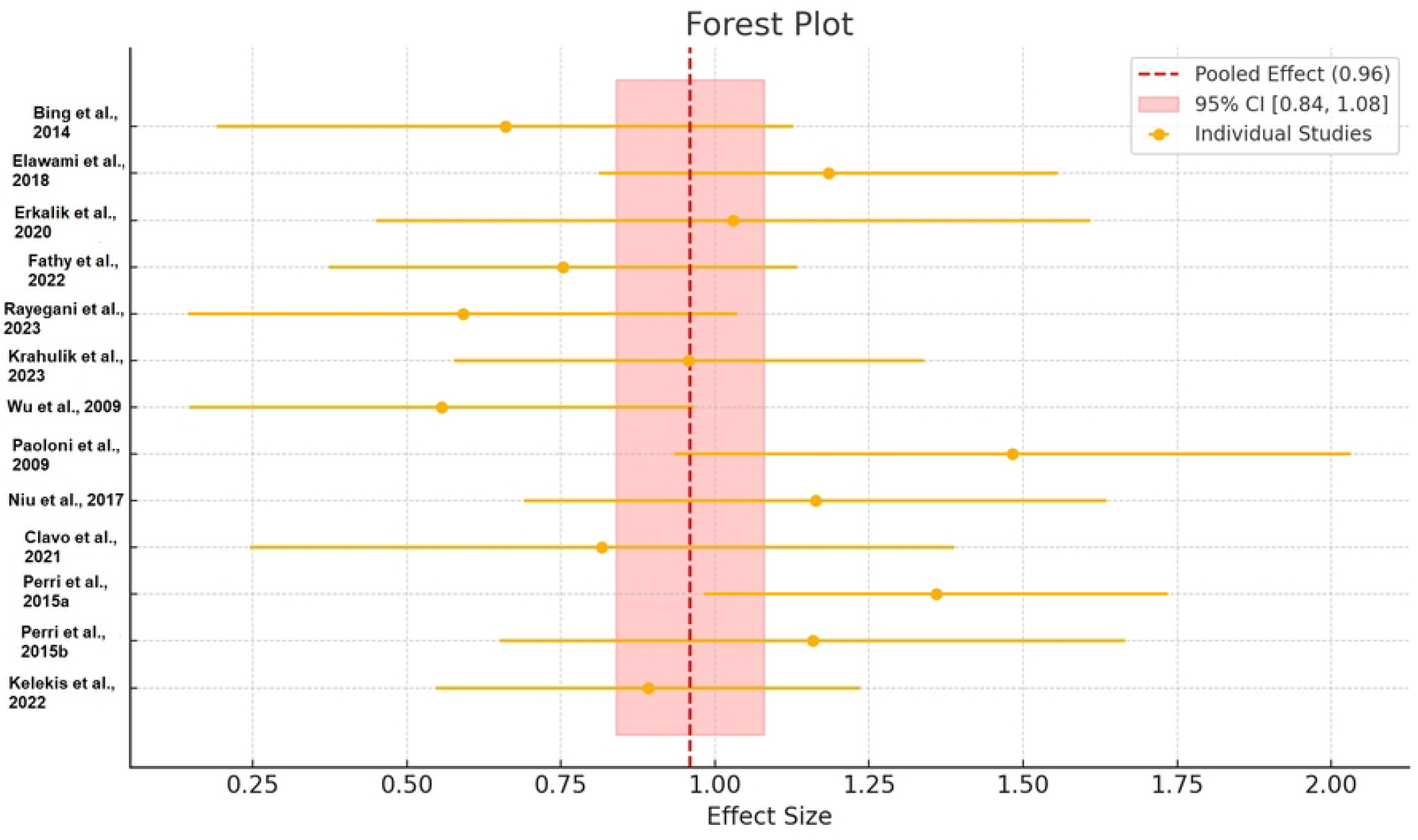
This forest plot visualizes the effect sizes from individual studies, along with the pooled effect size and its 95% confidence interval (CI), assessing the overall impact of oxygen-ozone therapy on spine disorders. Each yellow point represents the effect size from a specific study, and the horizontal line indicates the 95% confidence interval for that study. The studies are labelled on the y-axis (e.g., “Bing et al., 2014”), providing references to the source of the data. Wide horizontal lines indicate greater variability or lower precision in the individual study results. The red dashed vertical line represents the pooled effect size (0.96), calculated as the weighted average of the individual study effect sizes. The weighting considers the precision (inverse variance) of each study. The pink shaded region spans the pooled 95% CI [0.84, 1.08], indicating the range in which the true effect size is expected to lie with 95% certainty. This confidence interval helps assess the consistency and significance of the pooled estimate. Most individual study points cluster near the pooled effect size of 0.96, suggesting general agreement across studies. Some studies, such as “Wu et al., 2009,” have wider CIs, likely due to smaller sample sizes or methodological variability. The overall pooled effect suggests a positive therapeutic outcome, with the CI encompassing 1.0, indicating potential clinical significance. Plotted with Python *matplotlib* and *seaborn* for plotting. Data elaboration tools like *numpy* and *pandas* for meta-analysis calculations were adopted.

**Figure 9.**
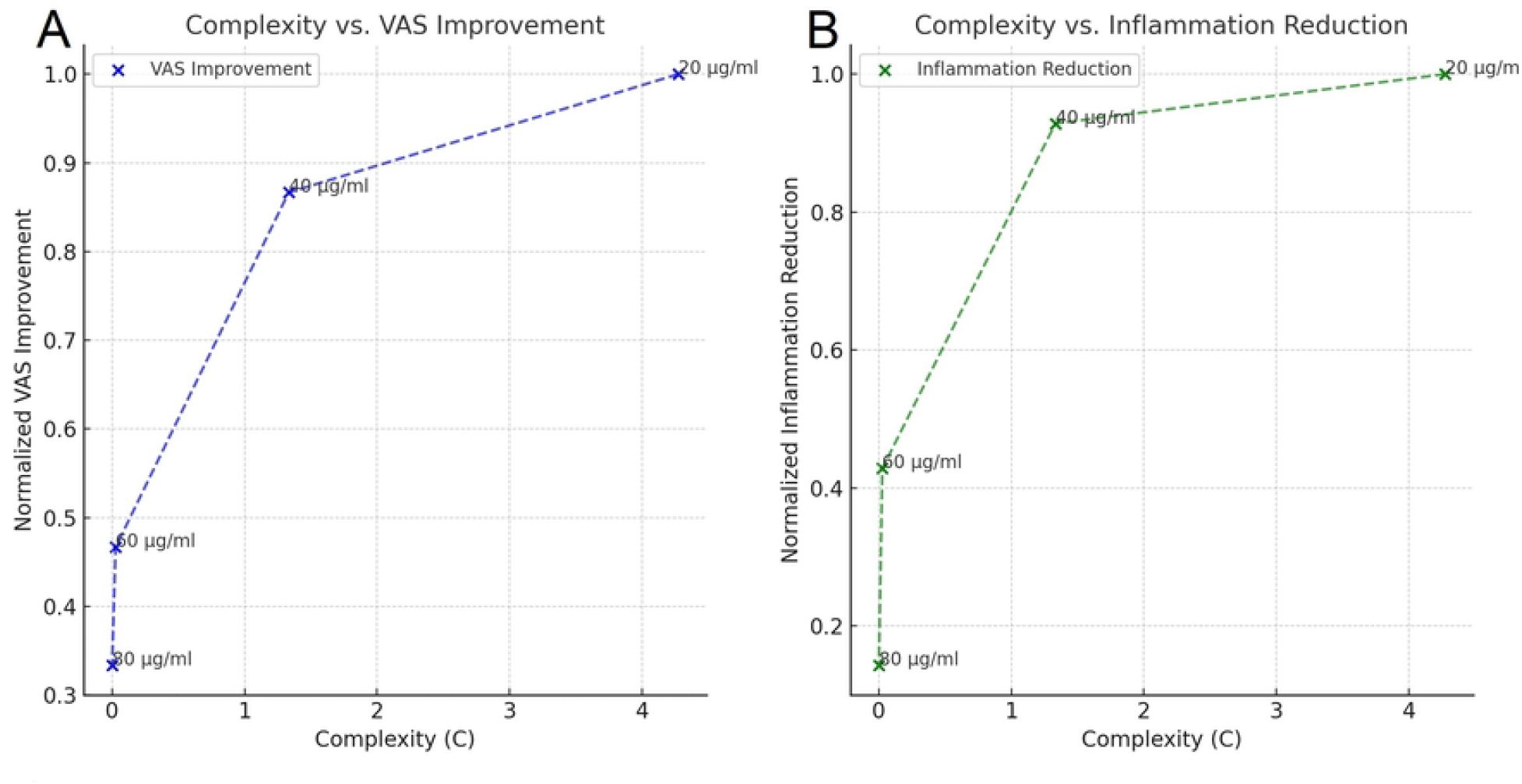
The plot shows a positive relationship between system complexity (C) and Visual Analog Scale (VAS) improvement. As ozone dose decreases from 60 µg/ml O_3_ to 20 µg/ml O_3_, complexity rises sharply, corresponding to enhanced VAS improvement. The most significant improvement is observed at 20 µg/ml O_3_, where complexity reaches a maximum (~4), normalized VAS improvement approaches 1.0, and system recovery is optimized. Similarly, a positive correlation exists between complexity and inflammation reduction. Higher complexity (C) achieved with lower ozone doses (20–40 µg/ml O_3_) corresponds to improved inflammatory control. At 20 µg/ml O_3_, the system reaches the lowest inflammation levels, reflecting an optimal therapeutic effect. Complexity is derived using metrics such as Shannon entropy and fractal dimensions to capture adaptive system behaviour. Normalized VAS and inflammation data were mapped against complexity to identify trends. Python *matplotlib* for visualization, with *numpy* handling data normalization. Plotted with Python code (Python 3.9) in Matlab (R2022a) environment.

Complexity vs VAS reduction (Figure 9A, r = 0.88, p = 0.12) and complexity vs inflammation reduction (Figure 9B, r =0.81, p = 0.19), show strong positive correlation but without statistical significance, probably due to the current data size.

The peaks at 20 μg/ml O_3_ suggest that this dose optimally balances the system’s complexity to maximize both pain reduction and inflammation control in spine musculo-skeletal disorders. Moreover, the steep improvements from 60 μg/ml O_3_ to 40 μg/ml O_3_ and then to 20 μg/ml O_3_ reflect the increasing ability of the system to adapt and stabilize. Higher doses (80 μg/ml O_3_ and 60 μg/ml O_3_) fail to induce sufficient complexity, resulting in reduced therapeutic outcomes.

Figure 10 shows how the different doses behave along time (minutes) respect to disorder (Shannon entropy), complexity and induction or modulation of chaos. The overall picture reported in the plot cannot allow to distinguish a clear difference among different doses. Entropy (Figure 10A) remains fluctuating across all doses but varies in amplitude, suggesting that ozone affects the system’s randomness similarly across doses. Minor variations between doses might indicate different impacts on system organization. As regarding chaos (Figure 10B), fluctuations in λ highlight dose-dependent chaos and stability. Higher doses (e.g., 80 µg/ml O_3_) may show larger fluctuations, representing higher adaptive stability or chaotic dynamics. Complexity (Figure 10C) fluctuates across all doses, reflecting the interplay between entropy and Lyapunov dynamics. The optimal dose for achieving adaptive chaos (high *C*) can be inferred by comparing the peaks and overall stability of *C* across doses.

**Figure 10.**
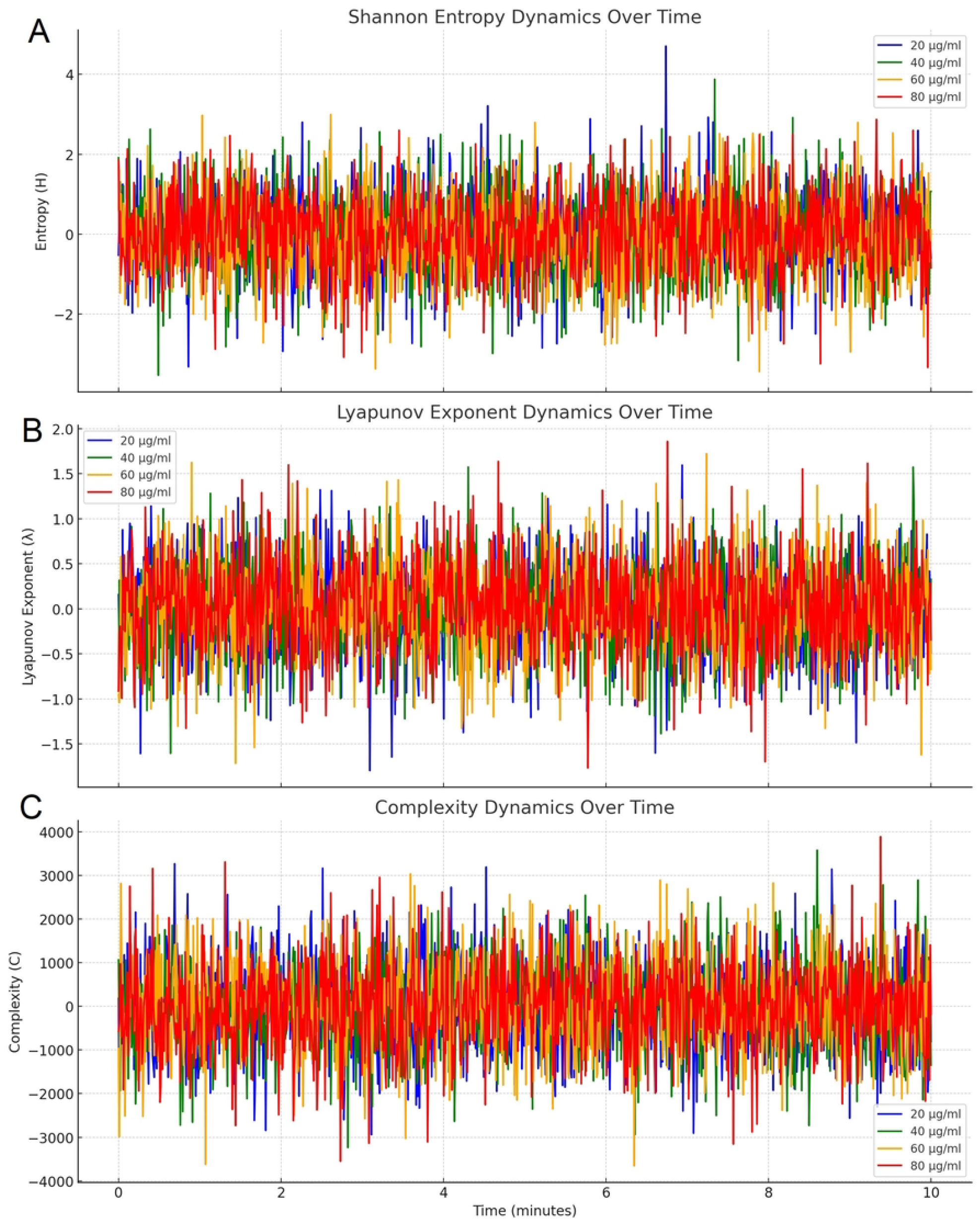
In Figure 10A, fluctuating entropy levels are depicted for four ozone doses (20, 40, 60, 80 µg/ml O_3_). Lower doses (20 and 40 µg/ml O_3_) exhibit smaller amplitude fluctuations, indicating controlled system entropy, while higher doses (60 and 80 µg/ml O_3_) display greater instability. In Figure 10B, Lyapunov exponent dynamics show dose-dependent chaotic behaviour. Lower doses (20 and 40 µg/ml O_3_) maintain adaptive chaos with lower peaks and minimal divergence, while higher doses reflect more turbulence. In Figure 10C, complexity dynamics stabilize at lower doses, while higher doses (60 and 80 µg/ml O_3_) lead to chaotic system behaviour with lower complexity values. Shannon entropy, Lyapunov exponent, and complexity were computed using time-dependent equations, incorporating Gaussian noise for biological variability. Plots were generated in Python with real-time data simulated over a 10-minute interval. Plotted with Python code (Python 3.9) in Matlab (R2022a) environment.

Paradoxically, all doses elicit dynamic responses, but the subtle differences in *S*, λ, and *C* suggest nuanced effects of each dose on system behaviour. This should suggest that the ability of a defied dose to promote the adaptive chaos depends on the level of chaoticity and Shannon entropy of the ozone-perturbed system, as adaptive chaos is elicited by an interplay between ozone dose and system’s dynamics. Actually, complexity dynamics (*C*) appear to indicate the ability of each dose to promote adaptive chaos, as it represents a balance between stability and flexibility, beneficial for biological systems.

The system’s baseline state (rigid, intermediate, chaotic) may influence how doses impact *S, λ*, and *C*.

Figure 11 helps us in addressing this apparent bias by evaluating ACM.

**Figure 11.**
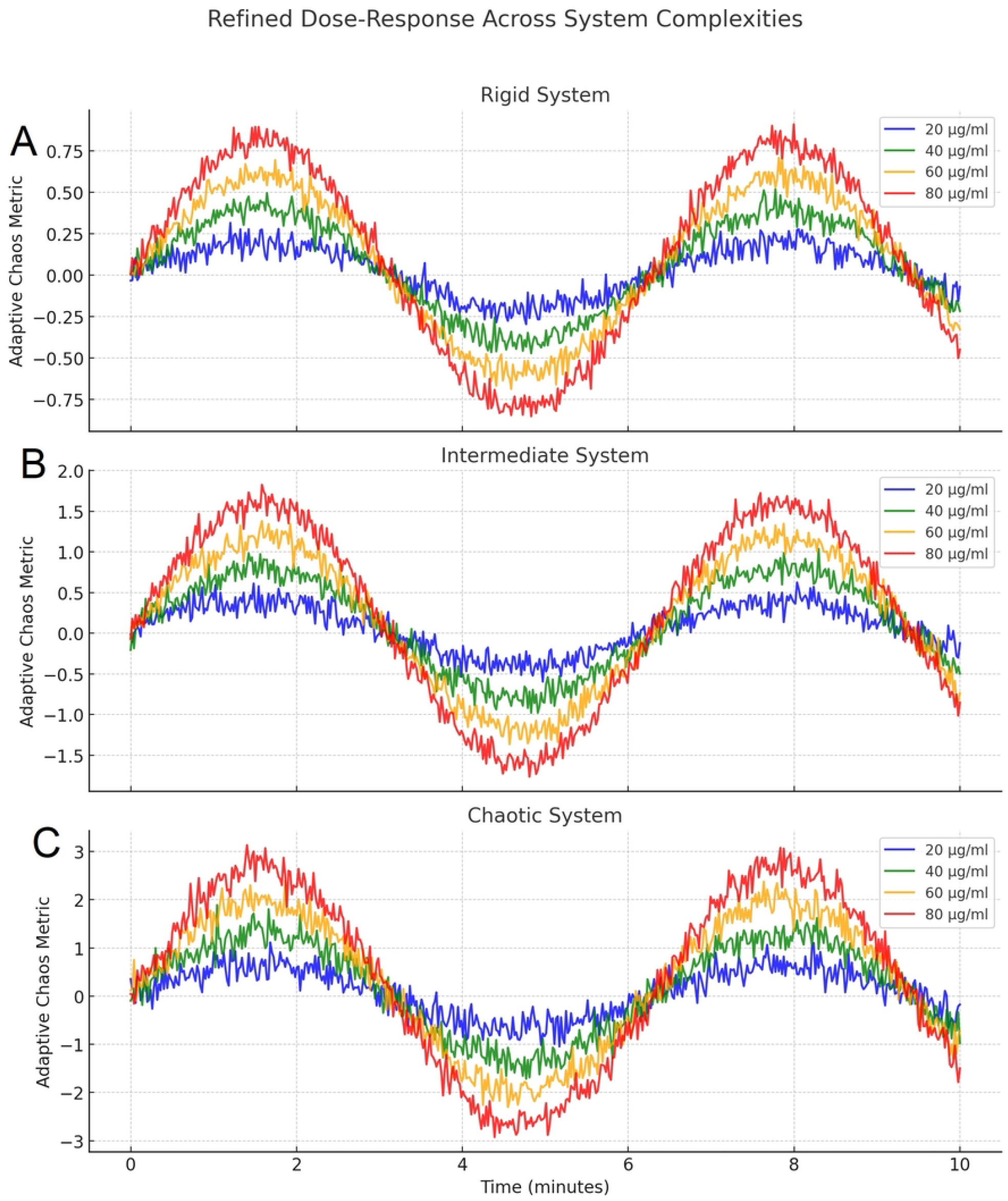
In Figure 11A (Rigid system) Adaptive Chaos Metric (ACM) is minimal across all doses due to low system flexibility, with higher doses (60 and 80 µg/ml O_3_) slightly outperforming lower doses. In Figure 11B intermediate systems show a clear dose-response relationship, with 20 µg/ml O_3_ yielding the highest ACM. This represents a “sweet spot” for achieving adaptive chaos. Figure 11C shows that in chaotic systems, lower doses (20 and 40 µg/ml O_3_) stabilize dynamics, while higher doses (60 and 80 µg/ml O_3_) amplify chaos. ACM was calculated by integrating Shannon entropy, Lyapunov exponents, and complexity. Dose responses were modelled with sinusoidal functions, incorporating baseline complexity. Plotted with Python code (Python 3.9) in Matlab (R2022a) environment.

While Figure 10 reports that all ozone doses have comparable variations without strong dose-specific separation, due to the strong impact of biological variability, so demonstrating system dynamics across metrics without a clear differentiation in adaptive chaos behaviour, Figure 11 focuses onto adaptive chaos, in a refined adaptive chaos metric across system types. It represents adaptive chaos metrics for three system types: rigid (Figure 11A), intermediate (Figure 11B), and chaotic (Figure 11C), demonstrating clear dose-dependent responses.

In Figure 11A rigid systems exhibit low flexibility and high order, are resistant to perturbations and have difficulty adapting to new states. They perform low baseline complexity and minimal response to ozone doses, as seen in Figure 11A, where adaptive chaos metrics are suppressed. Here, ozone doses induce small perturbations but fail to elicit significant adaptive chaos. Higher doses (80 µg/ml O_3_) tend to have slightly stronger effects, but the overall range of dynamics remains narrow.

Intermediate systems (Figure 11B) are moderately complex, with a balance between order and adaptability. They are more responsive to controlled perturbations. Figure 11B demonstrates more pronounced adaptive chaos metrics, with distinct separation between doses.

The doses 20 µg/ml O_3_ (blue) and 40 µg/ml O_3_ (green) elicit the most balanced adaptive responses. Intermediate systems are the “sweet spot” for ozone therapy, where the system can achieve significant adaptive chaos without tipping into excessive chaos or rigidity.

Chaotic systems are highly complex systems with inherent disorder. They exhibit high adaptability but may lack stability. Figure 11C shows strong dose responses, with all doses eliciting adaptive chaos. However, 20 µg/ml O_3_ and 40 µg/ml O_3_ maintain better control, while higher doses (60 µg/ml O_3_ and 80 µg/ml O_3_) amplify chaotic dynamics further. Chaotic systems are highly sensitive to ozone doses. Lower doses help stabilize the system, while higher doses exacerbate chaotic behaviour.

### Is it possible to forecast the best dose range on the basis of its adaptive chaotic features?

Forecasting the best dose of ozone to be used in clinics, even in the exemplificative circumstance of spine musculo-skeletal, painful disorders due to disc herniation, should rely on sophisticated bio-informatic algorithms, which can be even available in the next few years, standing on the current scientific progress in machine learning for medicine.

Figure 12 demonstrates the relationship between ozone dose (measured in µg/mL) and the adaptive chaos metric (ACM), showcasing how the adaptive chaos metric varies across different doses. A Gaussian model was employed to fit the observed data points and predict the dose that maximizes the adaptive chaos effect. The figure suggests that the adaptive chaos metric exhibits a bell-shaped dependency on ozone dose, with a peak at approximately 44.99 µg/mL O_3_. At this optimal dose, the system achieves the maximum adaptive chaos effect, a critical parameter for predicting therapeutic efficacy in applications utilizing ozone. The data indicate that at doses below or above the optimal value, the adaptive chaos metric decreases, suggesting a diminished effect. This behaviour highlights the importance of dose optimization to achieve the desired outcome in adaptive chaos-based interventions.

**Figure 12.**
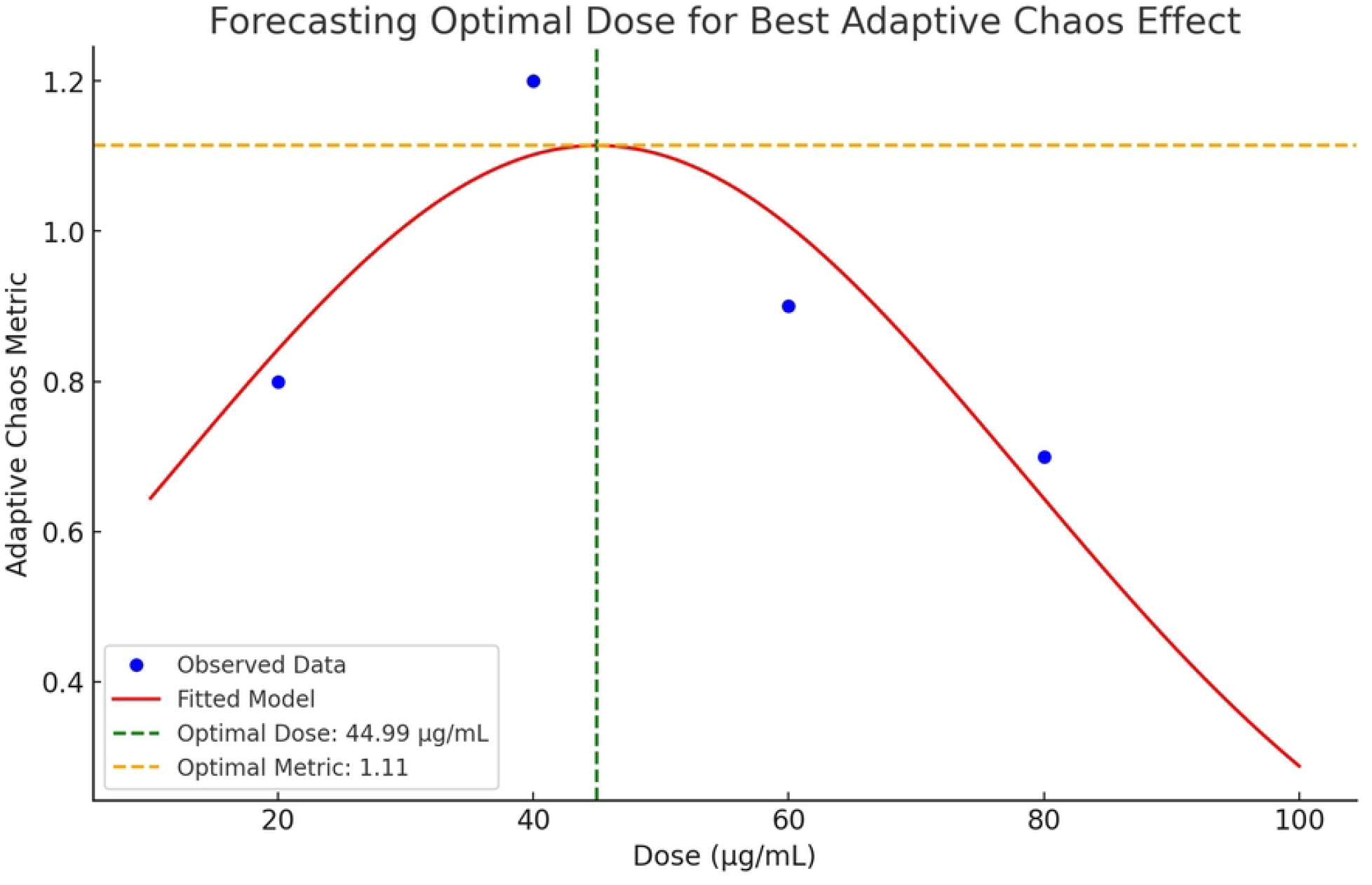
The graph predicts the optimal ozone dose for maximizing adaptive chaos. A Gaussian model (red curve) indicates the optimal dose at 44.99 µg/ml O_3_, with an Adaptive Chaos Metric (ACM) peak of 1.11. Observed data points (blue dots) align well with the fitted curve, validating the model. A Gaussian model was fitted to observed dose-ACM data using Python’s *scipy*.*optimize*.*curve_fit*. Adaptive chaos was predicted by balancing entropy, complexity, and chaos metrics. The red curve represents the Gaussian fit, and the green dashed line marks the optimal dose. Plotted with Python code (Python 3.9) in Matlab (R2022a) environment.

The study supports the hypothesis that an intermediate dose (44.99 µg/ml O_3_) maximizes the adaptive chaos effect, obviously, as outlined in this study, depending on the system state before ozone starting treatment. A dose of 45 µg/ml O_3_ is actually considered as the maximal possible to achieve the best clinical systemic efficacy (Figure 13), usually adopting major autohaemotherapy. This information is essential for therapeutic protocols that rely on precise dosing to optimize outcomes while minimizing undesired effects.

**Figure 13.**
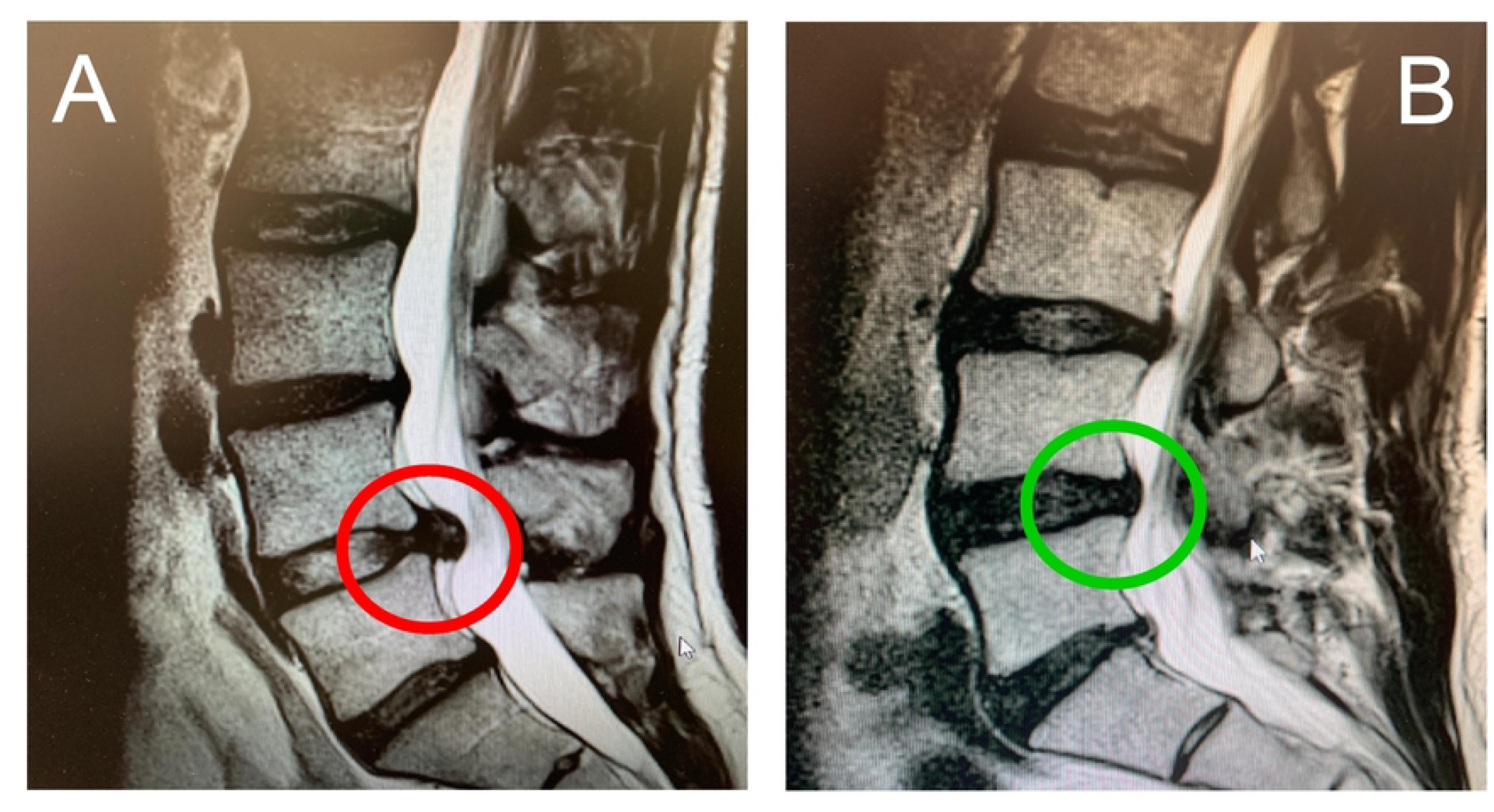
Sagittal Magnetic Resonance Imaging (MRI) scans of the lumbar spine, commonly used to assess spinal pathologies such as disc herniation, spinal stenosis, and degenerative changes, obtained in a male patient aged 66 years with chronic low back pain from June 2021. Figure 13A shows the vertebral bodies, intervertebral discs, spinal canal, and surrounding soft tissues before starting an oxygen-ozone therapy protocol. A notable feature in this image is the protrusion or herniation of an intervertebral disc into the spinal canal (red circle), possibly compressing the thecal sac or nerve roots. The spinal canal appears narrowed, suggesting spinal stenosis or discogenic compression. Figure 13B shows a sagittal lumbar spine view, following 65 days of oxygen-ozone therapy, assessing the disappearance of the herniation with protrusion and the complete disappearance of pain with improvement of motor disability. Oxygen-ozone therapy followed methods previously described with the inclusion of three session (1/week) of oxygen-ozone major autohaemotherapy (O_2_-O_3_-MAHT) at 45 µg/ml O_3_ as previously described. Images were obtained using a 3T MRI SIGNA™ scanner, T2-weighed sequences (3-5 mm for sagittal spine imaging) to provide high contrast, TR 4000 ms, echo time (TE) 120 ms, FOV 350 mm, matrix size 512×512. The patient signed an informed consent following criteria from Helsinki Declaration.

## Discussion

The evaluation of adaptive chaos in medical therapy is a highly innovative approach to measuring the evolution of biological dynamics as a whole, following treatment with a bioactive substance with curative properties^2,16,38,39^. Generally, phenomena governed by adaptive chaos behave according to a paradoxical pharmacokinetics, more broadly defined as hormesis^40^, and this is one of the reasons why our study considered oxygen-ozone therapy for painful musculoskeletal back disabilities as an illustrative model. Considering the idea that in the morpho-functional unit where inflammatory phenomena occur and are resolved, the control of chaos often occurs by reducing the complexity of the system, the parameters that measure chaos and complexity were used as functional markers of healing or health recovery. It should be noted that some concepts regarding the calculation of chaotic dynamics using Lyapunov exponents might appear contradictory, particularly when examining the lambda values for Figure 1 compared to Figure 2. This evidence summarizes the difficulty in focusing chaos parameters as their evaluations depend on the targeted system and the metrics adopted in a defined model.

Figure 1A shows exponential cytokine dynamics in Case A versus stable dynamics in Case B. This suggests an underlying deterministic system, as exponential growth is not characteristic of pure random noise. Figure 1B shows that Lyapunov exponents (λ) are exceptionally large, indicating high chaos in Case A and low chaos in Case B. This could reflect Gaussian noise rather than deterministic dynamics. Figure 1C reports that Shannon entropy rises sharply in Case A, correlating with cytokine dynamics and suggesting high randomness. Moreover, Figure 2A reports that Lyapunov exponents are smaller and realistic for deterministic systems, with negative λ (chaotic decay) for Case A and positive λ (adaptive chaos) for Case B. Figure 2B shows that complexity index is calculated, with Case B reporting extremely high values due to reduced Shannon entropy and structured functional relationships. Differences, particularly in λ values, exist because of the use of separate methods. For Figure 1 were used, as inputs, purely random time series with Gaussian noise added to model the scenario, whereas for Figure 2 were adopted deterministic ODE solutions with biological significance, modulated by noise. Discrepancies, therefore, arise because Figure 1 measures random processes, producing values that reflect noise intensity rather than deterministic chaos, whereas Figure 2 captures true chaos and complexity in modelled biological systems, providing more meaningful insights, close to the real world.

Data modelling in this study, ascertained that higher doses in the estimated hormetic range 20-80 µg/ml O_3_, exhibited a high Shannon entropy, whereas lower doses showed a greater ability in promoting complexity, with high fractal dimensions.

Figure 5 highlights, actually, the dose-dependent nature of fractal dimensions in relation to system adaptability: lower concentrations (20–40 µg/ml O_3_) promote adaptive chaos, leading to higher fractal dimensions and greater complexity over time. Higher concentrations (60–80 µg/ml O_3_) reduce adaptive capacity, limiting fractal dimension growth and reflecting disrupted system organization.

Figure 7 showed that the lowest dose exhibited the highest ability in inducing adaptive chaos. Low turbulence across the timeline indicates that 20 µg/ml O_3_ does not induce pathological states, unlike higher doses. This could be a marker for therapeutic efficacy. Furthermore, the coupling of chaos and complexity demonstrates the importance of dose optimization in therapeutic interventions. Future studies could explore how different doses shift the balance between chaos, complexity, and turbulence.

Actually, data from Figure 7 provide a novel framework for understanding the dynamics of ozone exposure in biological systems. It underscores the interplay between chaos, complexity, and turbulence, demonstrating how low doses can promote adaptive behaviour while avoiding destabilization. This analysis supports the hypothesis that adaptive chaos is dose-dependent, with optimal ozone concentrations (20–40 µg/ml O_3_) enhancing complexity, while higher doses suppress it, potentially leading to dysfunction.

The evidence reported in this study collectively highlights the nuanced dose-dependent dynamics of oxygen-ozone therapy in modulating systemic chaos and complexity. Key results show that lower ozone doses (20–40 µg/ml O_3_) foster adaptive chaos, marked by reduced Shannon entropy and increased complexity, as reflected in measures like Lyapunov exponents and fractal dimensions. These metrics suggest a balance between order and chaos, essential for biological stability and repair, particularly in spinal disorders^19^.

Shannon entropy decreases exponentially at lower doses, stabilizing system dynamics (Figures 3A and 3B). In contrast, higher doses (60–80 µg/ml O_3_) exhibit sustained elevated entropy, reflecting excessive oxidative stress and disordered chaos, which undermine therapeutic outcomes. Complexity metrics further support this, with lower doses demonstrating significant increases in complexity over time (Figure 4), indicative of enhanced system adaptability and resilience. Conversely, complexity remains suppressed at higher doses, highlighting reduced interconnectivity and impaired structural organization.

Furthermore, fractal dimensions provide additional insights into adaptive behaviour, with lower doses achieving higher fractal dimensions, indicative of improved systemic adaptability and structural coherence (Figure 5). This aligns with strong negative correlations between Shannon entropy and fractal dimension for lower doses, such as −0.90 for 20 µg/ml O_3_ and −0.83 for 40 µg/ml O_3_ (Figure 6). Higher doses, however, show weaker correlations, signaling diminished capacity for adaptive chaos.

Real-world oscillatory dynamics validate these findings, showing that 20 µg/ml O_3_ ozone promotes synchronized oscillations between chaos and stability, indicative of optimal therapeutic conditions^19^ (Figure 7). Additionally, clinical correlations demonstrate that lower doses correlate with greater reductions in pain (VAS improvement) and inflammation markers, with complexity positively linked to these outcomes (Figure 9). Finally, predictive modelling identifies an optimal dose of approximately 45 µg/ml, maximizing the adaptive chaos metric while maintaining system stability (Figure 12).

In summary, these results underscore the therapeutic potential of oxygen-ozone therapy at lower doses, which optimize adaptive chaos, enhance biological complexity, and facilitate recovery. Higher doses, by contrast, disrupt this balance, emphasizing the importance of precision dosing in clinical applications.

What emerges from these results, therefore, confirms previous considerations about the use of this adjunct therapeutical approach in medicine. In the hormetic range, represented in Figure 12, the lowest part of the hormetic curve includes doses able to promote adaptive chaos in turbulent and/or rigid, low complex micro-environments, via the reduction of high chaos and the increase in a complexity characterized by high fractality, i.e., much more ordered and functionally interrelated.

Conversely, the evidence reported in Figure 10, then elucidated in Figure 11, recalls the puzzling nature of biological variability, for which all the doses apparently behave with similar patterns at a first glance, unless they are compared with the finest texture of chaotic parameters occurring in much more narrow times.

Highlighting these parameters in order to apply them in a possible quantitative evaluation to scan patient’s health or forecasting the therapeutical success of certain doses for targeted patients, is yet challenging to date, despite such studies offer the opportunity to open a window towards a deeper comprehension of biological complexity.

In conclusion, adaptive chaos is a cornerstone concept in modern medicine, offering insights into the reparative mechanisms of biological systems. By reframing mild chaotic phenomena as potentially beneficial, this paradigm enriches our understanding of health and disease. It advocates for therapies that enhance dynamic equilibrium, emphasizing the preservation of biological complexity as essential for effective medical care. The present study pioneers a multi-dimensional analysis of the impact of oxygen-ozone therapy on spinal disorders, bridging mathematical modelling with clinical evidence to optimize therapeutic strategies. This work not only deepens our grasp of ozone role in modulating chaos but also sets the stage for broader applications of adaptive chaos in personalized and systemic medicine.

### Limitations of the study

The study presents significant advancements in understanding adaptive chaos and its potential application in oxygen-ozone therapy for spinal disorders, as an exemplificative scenario of the use of this therapeutical approach. However, it also faces critical limitations. Firstly, the reliance on mathematical modelling and bioinformatic tools introduces a layer of abstraction that may not fully capture the complexity of biological systems. While metrics like Shannon entropy and Lyapunov exponents offer innovative ways to quantify therapeutic outcomes, they are heavily dependent on assumptions that might not align with real-world biological variability. The use of simplified scenarios, as outlined in the modelling sections, reduces the intricacy of systemic interactions, such as cellular networks and tissue dynamics, potentially underestimating or oversimplifying critical variables.

Another limitation is the high variability in key parameters, such as cytokine concentrations and fractal dimensions, which are influenced by individual biological differences and environmental factors. This variability is exacerbated by the introduction of Gaussian noise into the models, which, while attempting to simulate biological unpredictability, might distort the reliability of outcomes. Furthermore, the unusually high Lyapunov exponents in some scenarios suggest potential inaccuracies in modelling chaotic behaviour, as noted in the discrepancy between Cases A and B. The reliance on retrospective fitting of the model to real-world data, drawn from selected studies, also highlights a potential bias, given the moderate heterogeneity in the included clinical evidence.

The study focus on specific ozone doses within a hormetic range neglects the broader context of patient-specific responses and systemic variability, limiting its translational applicability. Finally, the lack of direct experimental validation of the modelled outcomes restricts the study capacity to bridge theoretical insights with clinical practice, making the proposed dose-response relationship and therapeutic strategies more theoretical than practical at this stage. These limitations underscore the need for further experimental validation and a more nuanced integration of systemic biological complexities.

## Conclusions and future remarks

The study presents a ground-breaking approach to understanding oxygen-ozone therapy in the context of spinal musculoskeletal disorders, leveraging the innovative paradigm of adaptive chaos. By utilizing mathematical modelling and bioinformatic tools, the research establishes a novel framework to measure and modulate complexity within biological systems. Key findings demonstrate that oxygen-ozone therapy can optimize adaptive chaos at lower doses (20–40 µg/ml O_3_) for therapy in spine disorders, effectively reducing inflammatory cytokines, enhancing systemic stability, and promoting structural repair in intervertebral disc degeneration. This contrasts with higher doses, which induce excessive oxidative stress and impair recovery, highlighting the importance of precise dosing.

A major advantage of this study lies in its application of metrics such as Shannon entropy, Lyapunov exponents, and fractal dimensions to quantify therapeutic outcomes and improve our knowledge of biological complexity. Actually, these metrics provide a multi-dimensional perspective on biological complexity, allowing researchers to determine optimal therapeutic windows with unprecedented accuracy. Furthermore, the integration of modelled insights with clinical evidence bridges theoretical constructs with practical medical applications, emphasizing the translational potential of these findings.

The clinical relevance of this study is underscored by its focus on precision medicine, advocating for personalized ozone dosing tailored to patient-specific dynamics. This represents a significant departure from traditional linear therapeutic approaches, introducing a systemic perspective that prioritizes resilience and adaptability in biological systems. The research also establishes a framework for extending adaptive chaos concepts to other domains of regenerative medicine, suggesting broad applicability beyond spinal disorders.

The novelty of this work lies not only in its scientific insights but also in its methodological rigor. The emphasis on controlled perturbations as a therapeutic strategy paves the way for interventions that enhance systemic complexity, offering a shift from merely suppressing symptoms to promoting systemic equilibrium. This paradigm challenges conventional medical approaches and positions adaptive chaos as a cornerstone for innovative therapeutic strategies in modern medicine.

## Notes

The Authors state they have no conflicts of interest

### Competing Interest Statement

The authors have declared no competing interest.

